# Functional and Taxonomic Diversity of Avian Communities Across Land-Use Gradients in Wayanad Using eBird Data

**DOI:** 10.1101/2025.05.31.657139

**Authors:** MP Sivachand, P Megha

**Affiliations:** School of Biology, IISER-TVM; School of Data Science, IISER-TVM

**Keywords:** eBird, Citizen Science, Aves, Wayanad, Western Ghats, LULC

## Abstract

Understanding bird diversity across land use and land cover (LULC) in fragmented landscapes is essential for effective conservation planning in human-modified landscapes. Using one year of eBird data from 2023, this study assessed bird species richness, diversity indices, and feeding guild composition across ten LULC classes in Wayanad, southern India. We employed rarefaction methods to estimate effort-adjusted species richness due to unequal sampling effort across classes, and analysed community composition using Jaccard dissimilarity and Principal Coordinates Analysis (PCoA). Natural habitats, primarily evergreen and moist deciduous forests, supported the most diverse and functionally specialised bird communities. In contrast, agricultural and urban areas were dominated by generalist guilds such as omnivores and granivores, indicating ecological simplification. Interestingly, tea plantations exhibited high rarefied richness despite lower overall diversity, likely due to uneven sampling patterns. The study demonstrates that citizen science data can reveal meaningful patterns in bird community structure across varied land use types with appropriate analytical corrections. These findings highlight the potential of eBird for landscape-scale biodiversity monitoring and support the value of integrating functional guild perspectives into conservation strategies.

## 1 Introduction

Wayanad district lies within northern Kerala, India’s ecologically significant Western Ghats. Covering approximately 2,131 km^2^, the district exhibits substantial topographic variation. Elevations range from about 700 meters in the eastern plains to 2,100 meters in the western highlands, supporting diverse habitats (Das, 2024). The high-altitude Shola habitats in the Western Ghats gradually slope eastward to form the Wayanad Plateau, which merges into the Deccan Plateau. This transitional landscape encompasses montane forests and grasslands, mid-elevation evergreen forests, riparian zones, wetland marshes, and moist deciduous forests (Easa & Rajesh, 2025). The Wayanad Plateau is a notable geomorphic feature within the otherwise dissected terrain of the Western Ghats (Sundarajan *et al*., 2014). The diverse landscape of Wayanad, interspersed with a mosaic of urban settlements and fragmented habitats, has created a complex environment for its fauna. Over the past century, the region has experienced significant landscape fragmentation. Large-scale inward migration and agricultural expansion during the mid-1900s led to the widespread conversion of forests into farmland, disrupting the continuity of habitats and breaking up expansive forest tracts into narrow, isolated patches (Nair *et al*., 1978). Human settlements and commercial plantations have increasingly separated the western evergreen forests from the eastern deciduous zones, intensifying habitat fragmentation (Anoop & Ganesh, 2020). Wayanad district supports a rich avian population comprising resident and migratory species, reflecting the region’s ecological richness. The district’s unique topography, varied microclimates, and a mosaic of habitat types influence the avian diversity. One notable species is the Banasura Chilappan (*Montecincla jerdoni*), listed as Endangered by the IUCN. Fewer than 2,000 individuals are estimated to survive, restricted to elevations above 1,200 meters in the mountainous shola forests of Wayanad. This species is a habitat specialist, highly adapted to dense shola vegetation (BirdLife International, 2024). Forested areas support such specialists, while agricultural and human-modified landscapes provide habitats for generalist species that often move between different habitats (Waltert *et al*., 2005). Endemic flora and fauna further enhance habitat quality, contributing to Wayanad’s significance as a key region for avian biodiversity conservation (Joseph, 2023). As of May 2025, citizen scientists and researchers have recorded over 360 bird species in Wayanad, based on 32,725 checklists submitted by 1,630 contributors to eBird, a citizen science platform managed by the Cornell Lab of Ornithology (eBird, 2025). Increasing landscape fragmentation poses significant challenges for bird communities by altering species composition, reducing abundance, disrupting movement patterns through limited resource access, and increasing edge effects. Understanding how avian diversity in such a fragmented landscape is crucial for effective conservation planning. This study aims to assess bird species diversity and habitat preference across different land-use types in Wayanad across seasons from the eBird-based data, thereby providing insights into citizen science-based data to see the land-use patterns of avian populations in this ecologically significant region.

## 2 Methodology

### Study Area

The study area, Wayanad district in Kerala, lies along the escarpment of the Western Ghats between N Lat. 11° 26^*′*^ 54^*′′*^ to 11° 58^*′*^ 52^*′′*^ and E Long. 75° 46^*′*^ 11^*′′*^ to 76° 25^*′*^ 8^*′′*^(Achu *et al*., 2021). Spanning 2,131 km^2^, approximately 40 % of Wayanad is under forest cover, administered by the North Wayanad, South Wayanad, and Wayanad Wildlife Divisions (Census of India 2011). The district is an east-sloping plateau with elevations ranging from 700–2100 m asl and drained by the River Kabini, one of the three east-flowing rivers in Kerala, a major tributary of the River Cauvery. It forms ecological continuums with Coorg in Karnataka to the north-west, the Nilgiris in Tamil Nadu to the southeast, and the Malabar plains to the west (Logan, 1887). The region receives an average annual rainfall of 2,322 mm with a distinct west-to-east rainfall gradient due to orographic effects (John *et al*., 2020). During the summer months (March to May), temperatures range between 20°C and 36°C, with reduced rainfall supporting the breeding activities of birds in grasslands and deciduous forests (Daniels, 1997). The southwest monsoon season (June to September) brings heavy rainfall, averaging 2500-3000 mm annually, revitalising rivers and dense evergreen forests but posing challenges like soil erosion and habitat disruptions (Pascal, 1988). This period supports rainforest species such as hornbills and kingfishers. From October to November, the northeast monsoon starts with low rainfall intensity. The Dry season (December to February) is marked by cooler temperatures, ranging from 15°C to 25°C, attracting migratory species while endemic birds like the Wynaad Laughing Thrush and Nilgiri Wood Pigeon thrive in shola forests and wetlands (Ali & Ripley, 1987).

**Figure 1.**
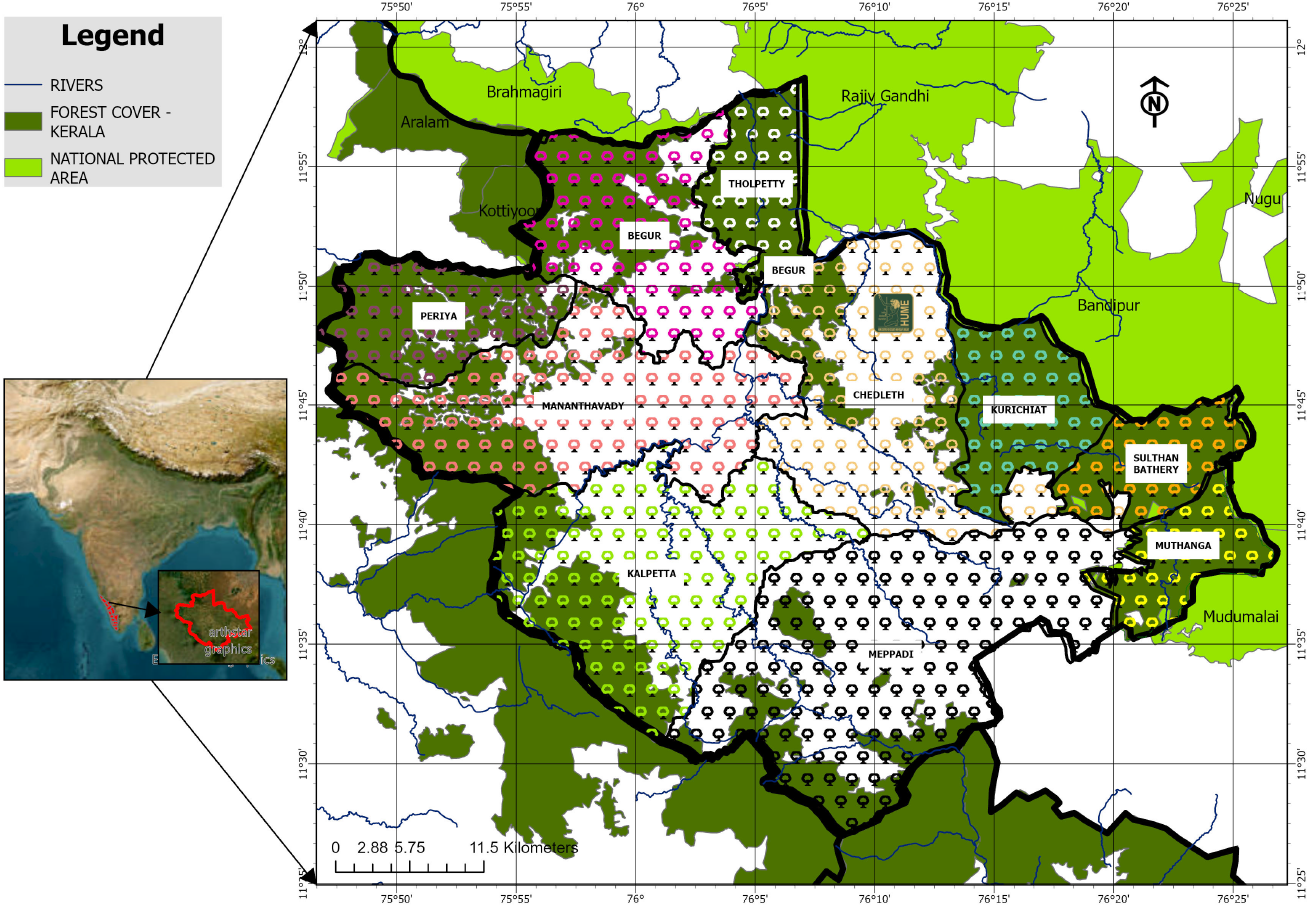
Map showing the study area.

### Data Collection

#### Collecting eBird data

eBird Open Data, managed by the Cornell Lab of Ornithology, is a globally accessible platform aggregating real-time bird observation data. This platform leverages contributions from citizen scientists, birdwatchers, researchers, and conservationists to document bird sightings across diverse species and locations. For this study, eBird data from January to December 2023 were utilised. The raw dataset underwent a meticulous cleaning to ensure its quality and reliability. Duplicates and unnecessary columns were removed to streamline the data for analysis. The cleaned dataset was then sorted based on the seasonality in Western Ghats, processed in ArcGIS Pro (Fig.2), where additional geographic information, such as elevation and land cover type, was extracted for each species.

**Figure 2.**
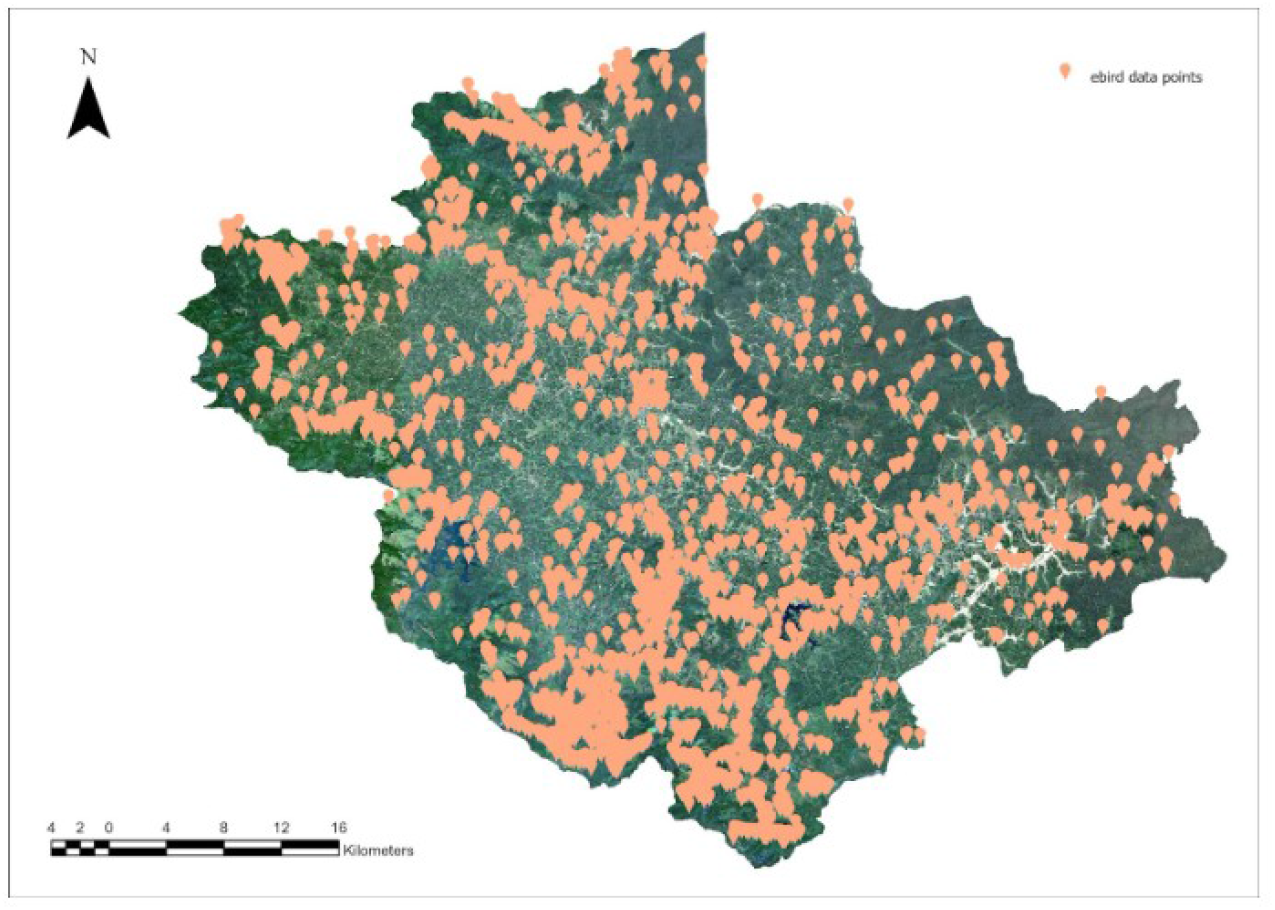
Map showing the distribution of eBird checklists submitted in Wayanad district, Kerala, India, from January 2023 to December 2023, illustrating spatial patterns of bird observation records.

#### Land Cover and Land Use Classification Using Sentinel-2 Imagery Image Acquisition

The study utilised Sentinel-2 imagery with a 10-meter spatial resolution to classify land cover and land use (LULC) patterns. Sentinel-2, equipped with a Multispectral Instrument (MSI) (European Space Agency [ESA], 2015), provides high-resolution imagery across multiple spectral bands, making it suitable for detailed LULC analysis. Specifically, Level-2A surface reflectance products were employed, which are atmospherically corrected and ready for analysis. Initially, the Sentinel-2 imagery for the study area and period of December 2023 was downloaded as .SAFE format from the Copernicus Open Access Hub. The selected tiles were preprocessed to ensure data quality. Preprocessing steps included cloud masking using the Sentinel-2 Quality Assessment Band (QA60) and spectral correction to minimise atmospheric distortions. These steps were performed using the Sentinel Application Platform (SNAP).

#### Preprocessing

Key spectral bands were extracted at a 10-meter resolution (e.g., Blue, Green, Red, and NearInfrared). Additional spectral indices, such as the Normalised Difference Vegetation Index (NDVI) and Normalised Difference Water Index (NDWI), were calculated to enhance feature differentiation. These indices facilitated the classification of vegetation, water bodies, and built-up areas. The study utilised supervised classification methods to categorise the LULC types. Training samples for distinct land cover classes, including built-up areas, vegetation, water bodies, and barren land, were manually delineated based on ground truth data, satellite imagery, and ancillary datasets. A machine learning algorithm, maximum likelihood classification (MLC), was implemented for classification. The algorithm was chosen for its robustness in handling high-dimensional data and its proven performance in remote sensing applications.

Normalised Difference Vegetation Index (NDVI)

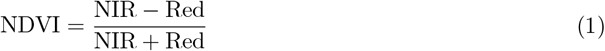

Normalised Difference Water Index (NDWI)

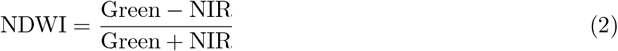

#### Post-classification

The results were validated using a confusion matrix and accuracy metrics. Validation was performed by comparing the classified results with independent ground truth data and high-resolution imagery. The confusion matrix was generated using the confusion matrix function from the scikit-learn library. This matrix provides a detailed breakdown of classification results. Accuracy, a measure of overall classification performance, was calculated as the ratio of correctly classified instances to the total number of predictions. This was computed using the accuracy score function from the same library. The formula used is:

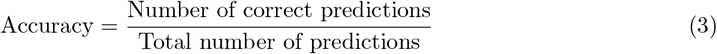

To enhance interpretability, the confusion matrix was visualised as a heatmap using the seaborn library.

#### Statistical Packages

All analyses were conducted using Python. Data manipulation was performed using pandas and numpy. Diversity and dissimilarity metrics were computed using the scikit-bio package. Guild composition, rarefaction, and ordination results were visualised using seaborn and matplotlib.

#### Guild Assignment

All bird species were assigned to one of six major feeding guilds based on published ecological traits: carnivores, frugivores, granivores, insectivores, nectarivores, and omnivores. Guild assignments were conducted at the species level using regional field guides and trait databases (Tobias *et al*., 2022).

Guild composition was then aggregated by LULC class, and the relative abundance of each guild was computed per habitat type.

#### Rarefaction and Diversity Indices

Rarefaction was used to standardise species richness estimates to address variation in sampling effort across LULC classes. Rarefied richness was calculated to a common number of observations per LULC class, allowing for unbiased comparisons. In addition to richness, we calculated Shannon and Simpson diversity indices to quantify species richness and evenness. These indices were computed separately for raw and rarefied datasets to assess the influence of sampling effort. The expected number of species in a rarefied sample of size *n* is calculated as:

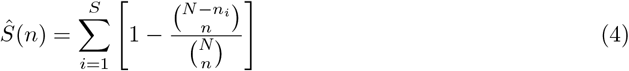

Where:

- *Ŝ*(*n*) is the expected number of species in a sample of size *n*,
- *N* is the total number of individuals across all species,
- *n*_*i*_ is the number of individuals of species *i*,
- *S* is the observed species richness,
- *n* is the rarefied subsample size.

Diversity was further quantified using the Shannon and Simpson indices. The Shannon Index is defined as:

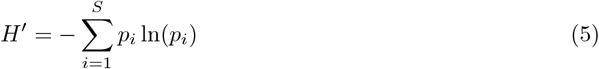

Where *p*_*i*_ is the relative abundance of species *i*. A higher *H*^*′*^ indicates greater species diversity and evenness.

The Simpson Index accounts for the probability that two individuals randomly selected from a sample belong to different species:

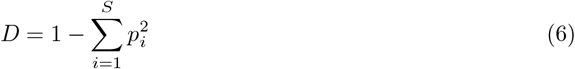

This index ranges from 0 to 1, where higher values denote greater diversity.

#### Dissimilarity and Community Ordination (PCoA)

We used the Bray-Curtis dissimilarity index to quantify differences in species composition among LULC classes. A Principal Coordinates Analysis (PCoA) was then conducted to visualise dissimilarity patterns in two-dimensional space. This method allowed us to detect similarities and differences in community structure across habitats, based on species abundances. In addition, a Jaccard similarity heatmap based on presence-absence data was generated to compare community overlap among habitats. To assess variation in bird community composition across LULC types, we used the Jaccard dissimilarity metric based on presence-absence data. The Jaccard Index between two sites is computed as:

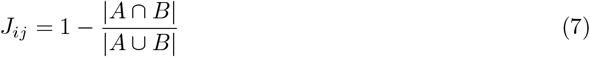

Where:

- *A* and *B* are the sets of species at sites *i* and *j* respectively,
- *|A ∩ B|* is the number of shared species,
- *|A ∪ B|* is the total number of unique species across both sites.

The resulting dissimilarity matrix was subjected to Principal Coordinates Analysis (PCoA) to visualize community structure in a reduced dimensional space. The PCoA is performed on a centered dissimilarity matrix **D**, and its eigen decomposition yields the principal coordinate axes:

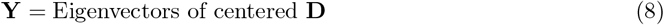

These eigenvectors represent sample scores in reduced dimensions and were used to identify clustering and compositional shifts among LULC classes.

## 3 Results

### Land Cover and Land Use Classification Using Sentinel-2 Imagery

The classification achieved an overall accuracy of 93.5 %, indicating strong performance. Higher accuracy was observed in distinct and larger classes, such as Mixed Crop – Dry Land and Evergreen Forest (95%). In contrast, visually similar or underrepresented classes like Wetlands and Rocks (80%) showed more misclassifications. These errors were primarily due to overlapping spectral features and imbalanced training data. Enhancing class separability and improving sampling can further strengthen classification reliability. There is a significant variation in land cover sizes in the study area (Fig.3). The largest class, Mixed crop- Dry land, covers 930 sq. km, making it the dominant land cover type. This is followed by Moist deciduous at 412 sq. km and Evergreen forest at 381 sq. km. These three classes collectively account for a significant portion of the total land area. Built-up areas occupy 68 sq. km, representing a moderate contribution to the total land area. This class is significantly smaller than the largest categories but larger than the smallest ones. The smallest classes, Rocks (4 sq. km), Wetlands (7 sq. km), and Water body (17 sq. km), make minimal contributions to the total area. The range of class sizes, from 4 sq. km to 930 sq. km, highlights the diversity of land cover types (Fig.3).

**Figure 3.**
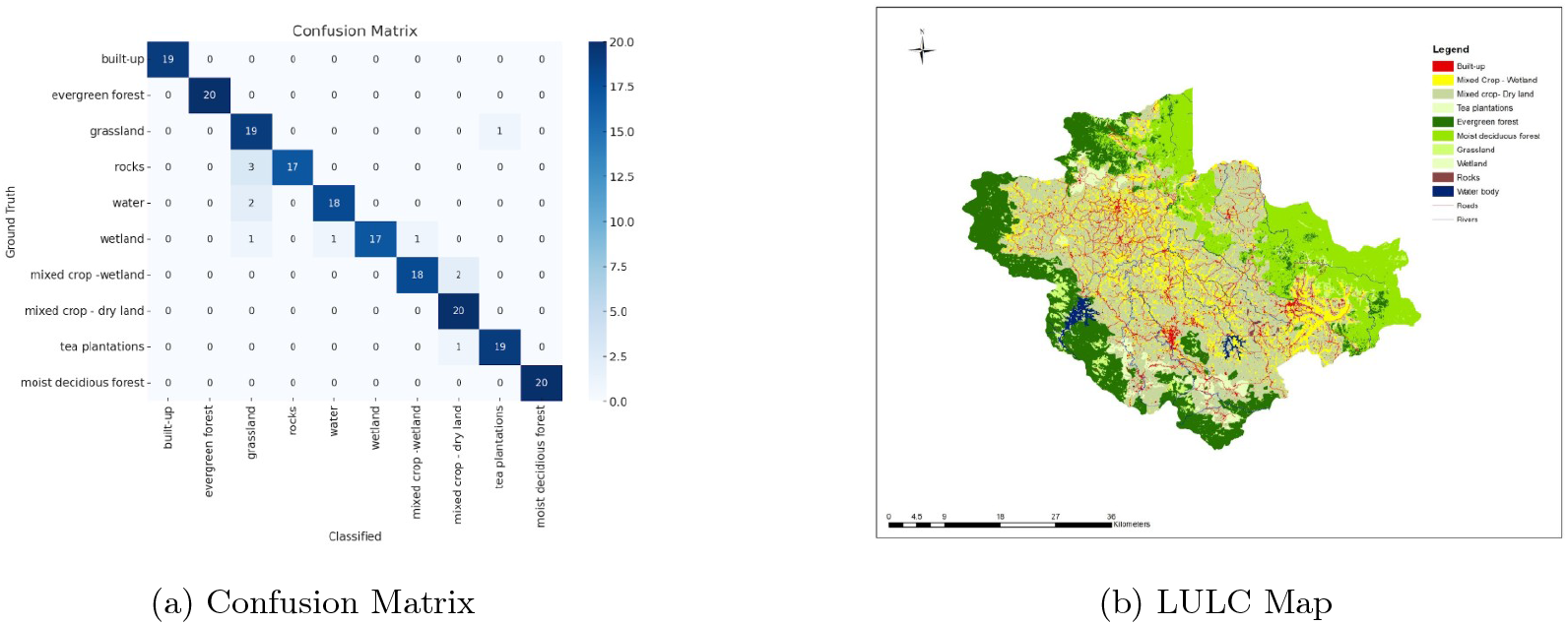
a. Confusion matrix evaluating the classification accuracy for each Land Use and Land Cover (LULC) class in Wayanad district, Kerala, India, derived from Sentinel-2 imagery analysis. b. Map displaying the Land Use and Land Cover (LULC) classification of Wayanad district, Kerala, India, derived from Sentinel-2 imagery.

**Figure 4.**
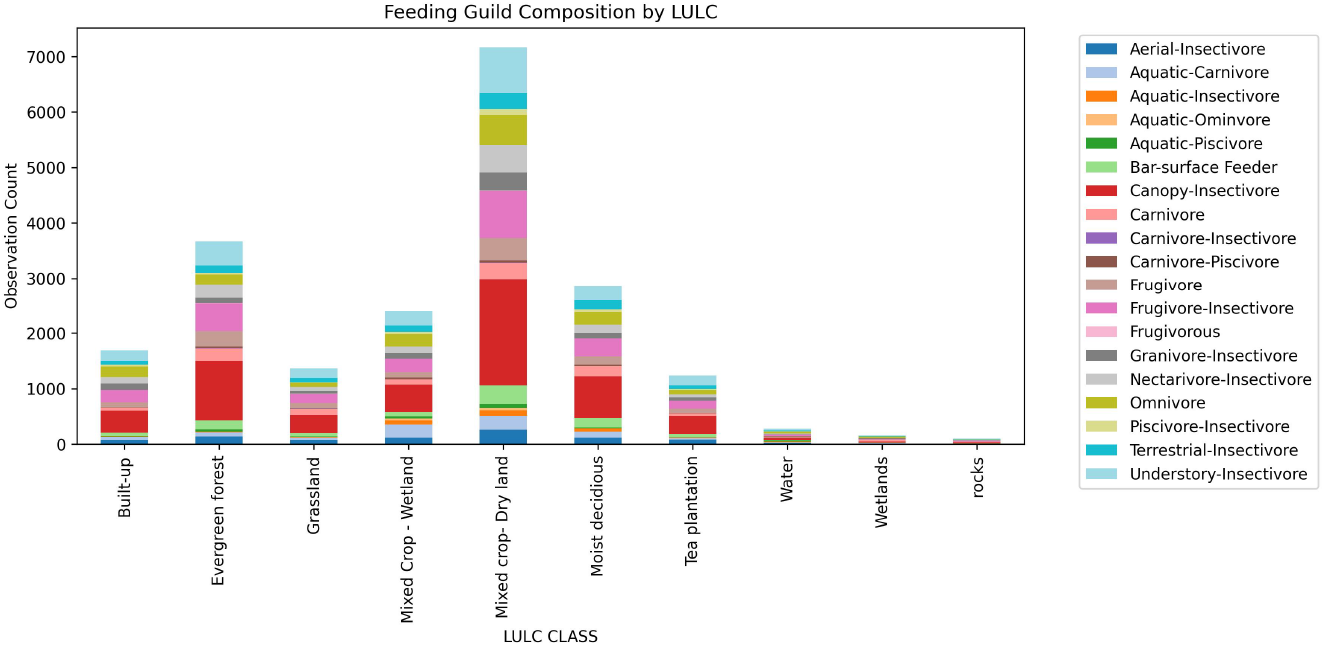
Feeding Guild Proportions across LULC Classes. This figure illustrates the proportional composition of feeding guilds in each land-use type, highlighting shifts in functional roles.

**Figure 5.**
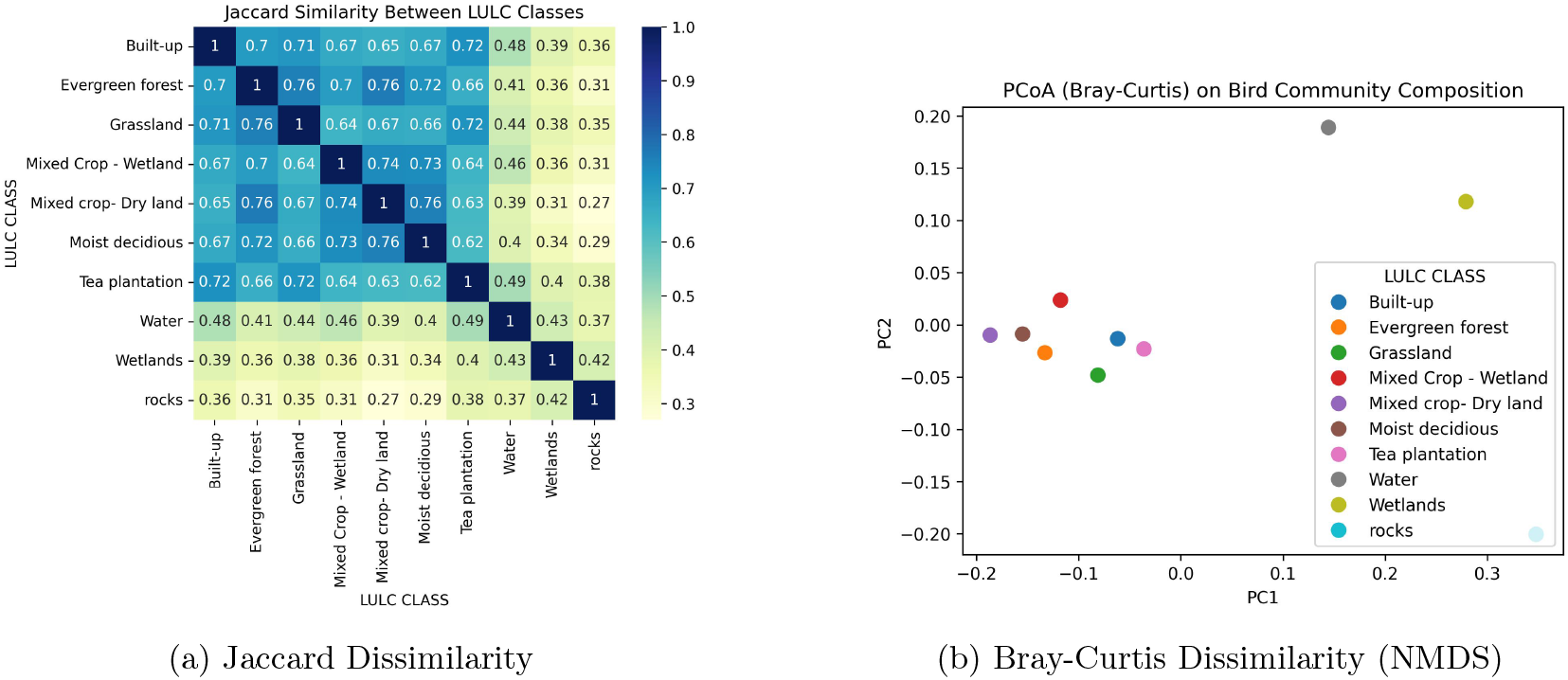
a. Jaccard Similarity Heatmap (Presence-Absence Based) across LULC Classes. This heatmap reveals pairwise species overlap among habitats, highlighting core communities and distinct assemblages. b. PCoA of Bray-Curtis Dissimilarity in Bird Community Composition across LULC Classes. The figure shows how bird communities cluster based on habitat, reflecting community-level habitat preferences.

#### Rarefied Richness vs Raw Richness

Raw species richness was highest in dryland crop, moist deciduous forest, and evergreen forest habitats. However, after applying rarefaction to correct for sampling effort, evergreen forest, dryland crop, and mixed dryland crop maintained high species richness, while fallow and rocky habitats dropped in rank, reflecting low diversity after effort correction.

**Table 1:**
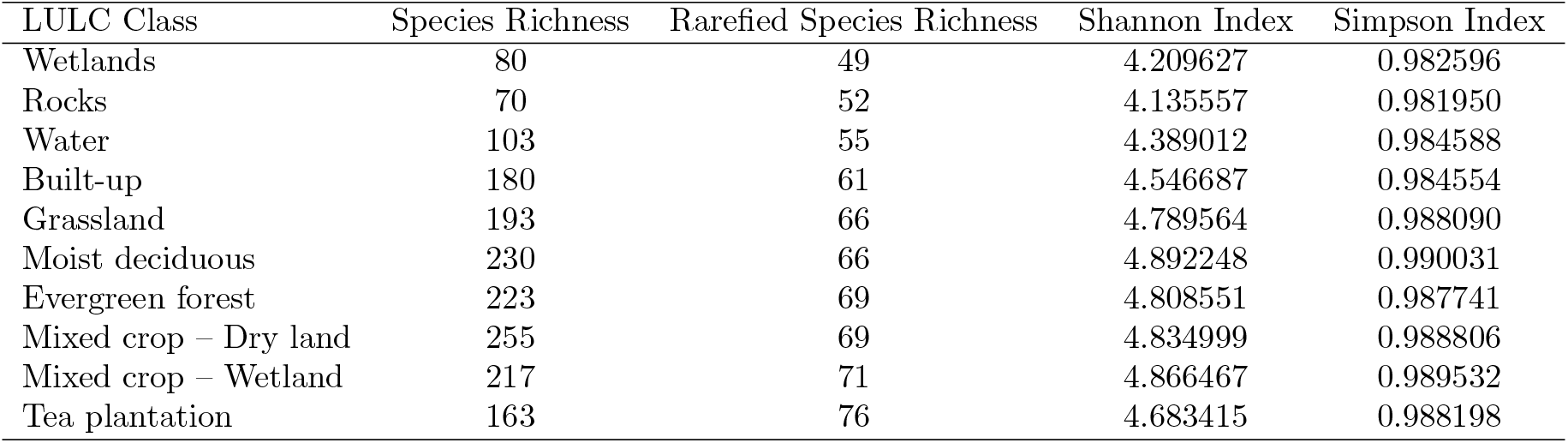
Rarefied vs Raw Species Richness across LULC Classes. This figure shows how rarefaction alters perceived richness, highlighting effort-independent diversity estimates.

| LULC Class | Species Richness | Rarefied Species Richness | Shannon Index | Simpson Index |
| --- | --- | --- | --- | --- |
| Wetlands | 80 | 49 | 4.209627 | 0.982596 |
| Rocks | 70 | 52 | 4.135557 | 0.981950 |
| Water | 103 | 55 | 4.389012 | 0.984588 |
| Built-up | 180 | 61 | 4.546687 | 0.984554 |
| Grassland | 193 | 66 | 4.789564 | 0.988090 |
| Moist deciduous | 230 | 66 | 4.892248 | 0.990031 |
| Evergreen forest | 223 | 69 | 4.808551 | 0.987741 |
| Mixed crop – Dry land | 255 | 69 | 4.834999 | 0.988806 |
| Mixed crop – Wetland | 217 | 71 | 4.866467 | 0.989532 |
| Tea plantation | 163 | 76 | 4.683415 | 0.988198 |

#### Feeding Guild Distributions

Feeding guild composition varied significantly across LULC types. Forest habitats (evergreen, moist deciduous) hosted more insectivores and frugivores, while granivores and omnivores dominated agricultural classes. Built-up areas had fewer specialised guilds and were primarily occupied by omnivores and generalists. Wetland and rocky areas showed limited guild diversity, often with dominance by a single group (e.g., carnivores in wetlands).

#### Community Composition (PCoA and Jaccard Index)

PCoA revealed apparent clustering by LULC class, with wetlands, rocks, and fallow areas forming distinct communities, while forests and agricultural areas exhibited overlapping composition. The first two axes of the PCoA explained a substantial portion of the variation in species composition, indicating ecological separation based on habitat type. The Jaccard similarity heatmap indicated that built-up, evergreen forest, and dryland crops shared high species overlap. In contrast, wetlands and rocky areas had low similarity with other classes, suggesting habitat-specific species pools.

## 4 Discussion

The analysis of avian communities across varied land use and land cover (LULC) classes reveals notable species richness, diversity, and functional guild distribution patterns. Natural forest habitats, particularly evergreen and moist deciduous forests, consistently supported a rich and balanced representation of specialised feeding guilds such as insectivores, frugivores, and carnivores. These patterns suggest structurally complex habitats retain high functional integrity and ecological niches. In contrast, human-altered landscapes, including mixed crop fields, tea plantations, and built-up areas, exhibit shifts toward generalist guilds like omnivores and granivores. This shift indicates reduced habitat specialisation and complexity, potentially lowering ecological resilience. Interestingly, tea plantations showed high rarefied species richness despite having lower overall observed richness and moderate diversity indices. This anomaly is likely due to fewer checklists with high internal species diversity, illustrating how rarefaction can overestimate richness in sparsely sampled but compositionally variable habitats. Such insights emphasise the need to interpret effort-standardised metrics alongside raw diversity measures. Despite inherent biases in eBird data, such as unequal sampling, observer variability, and lack of detectability estimates, using rarefaction and presence-absence metrics allowed us to uncover consistent ecological signals. Community composition analyses using Jaccard distances and PCoA further supported the differentiation of bird assemblages across LULC classes, highlighting distinct ecological communities even in habitats with low species counts, such as wetlands and rocky outcrops.

## 5 Conclusion

This study demonstrates the viability of using citizen science data, when properly filtered and statistically adjusted, to detect meaningful patterns in avian diversity and guild structure across landscapes with varying degrees of human modification, despite unequal sampling effort among LULC classes, rarefaction and community dissimilarity analyses revealed clear distinctions in species richness, functional guild representation, and habitat specificity. Natural forests serve as crucial reservoirs for diverse and specialised bird communities, while specific agricultural systems like mixed cropping fields may offer complementary habitat value. Conversely, simplified environments such as urban and monoculture plantations tend to favour generalist species, underscoring the ecological cost of land-use intensification. The rarefied richness anomaly in tea plantations further highlights the importance of combining multiple indices and interpreting them within their sampling context. When analysed with effort-aware tools, citizen science platforms like eBird can play a pivotal role in large-scale biodiversity monitoring and conservation planning. These findings support integrating participatory data into conservation strategies, especially in regions where conventional ecological monitoring is logistically challenging.

---

**Table 2:** Abbreviations of classes and seasons used in Table 3.

| LULC Classes |  |
| --- | --- |
| Built-up | BU |
| Evergreen forest | EVF |
| Grassland | GL |
| Mixed Crop - Wetland | MCW |
| Mixed crop - Dry land | MCD |
| Moist deciduous forest | MDF |
| Rocks | R |
| Tea plantation | TP |
| Water Body | WB |
| Wetlands | WL |
| Seasons |  |
| Dry season | DS |
| North east Monsoon | NEM |
| Summer | S |
| Southwest monsoon | SWM |

**Table 3:**
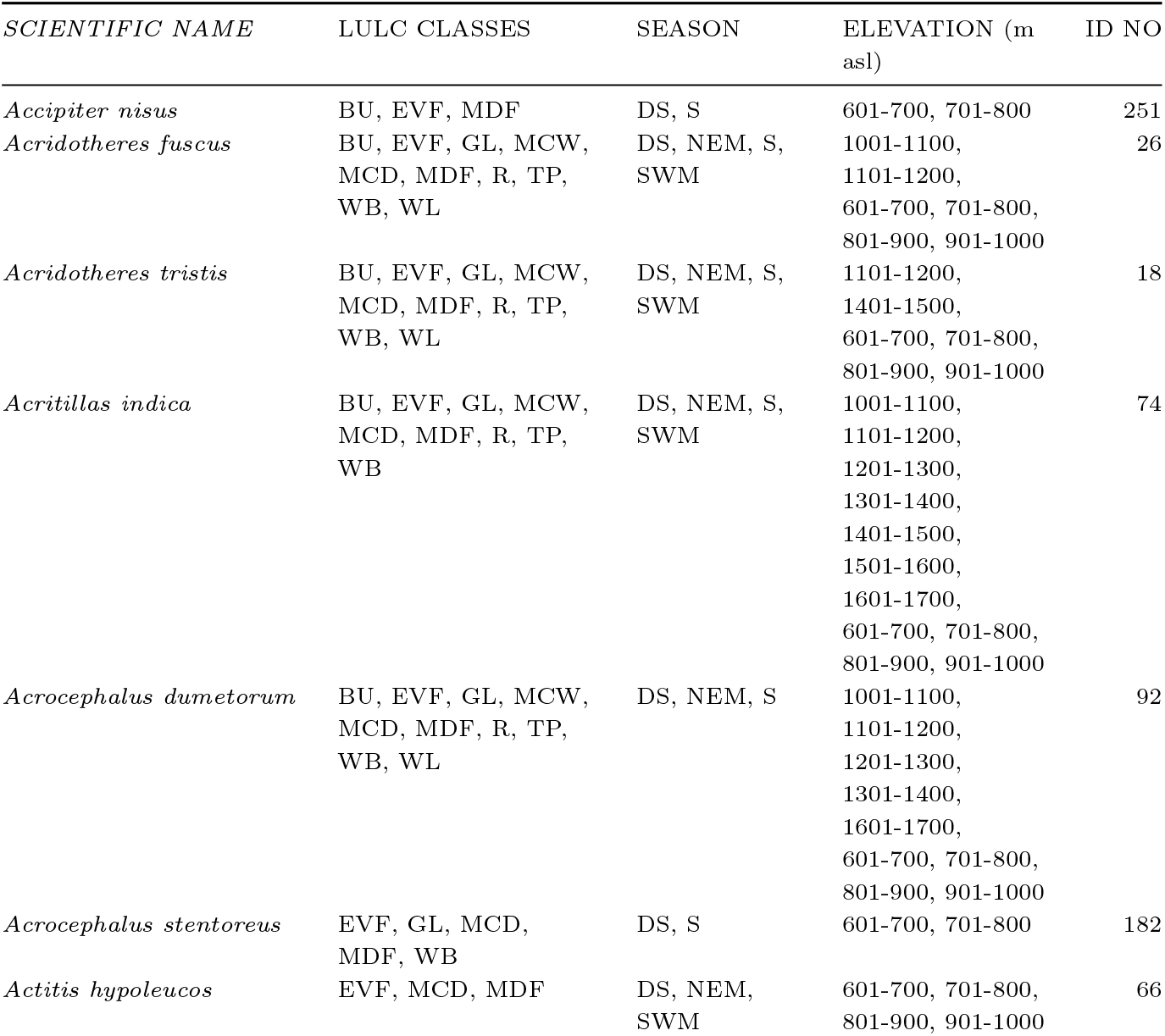
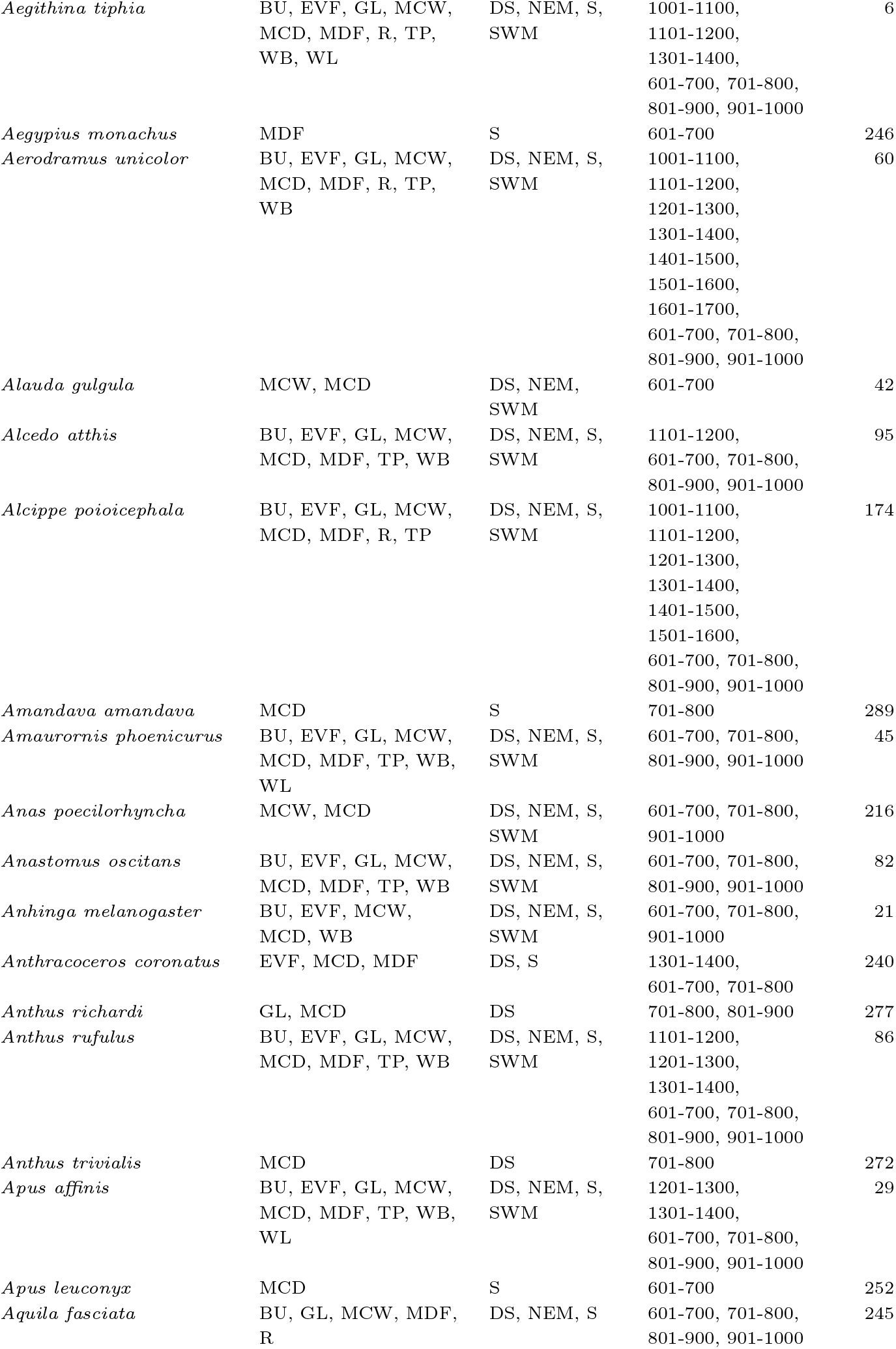
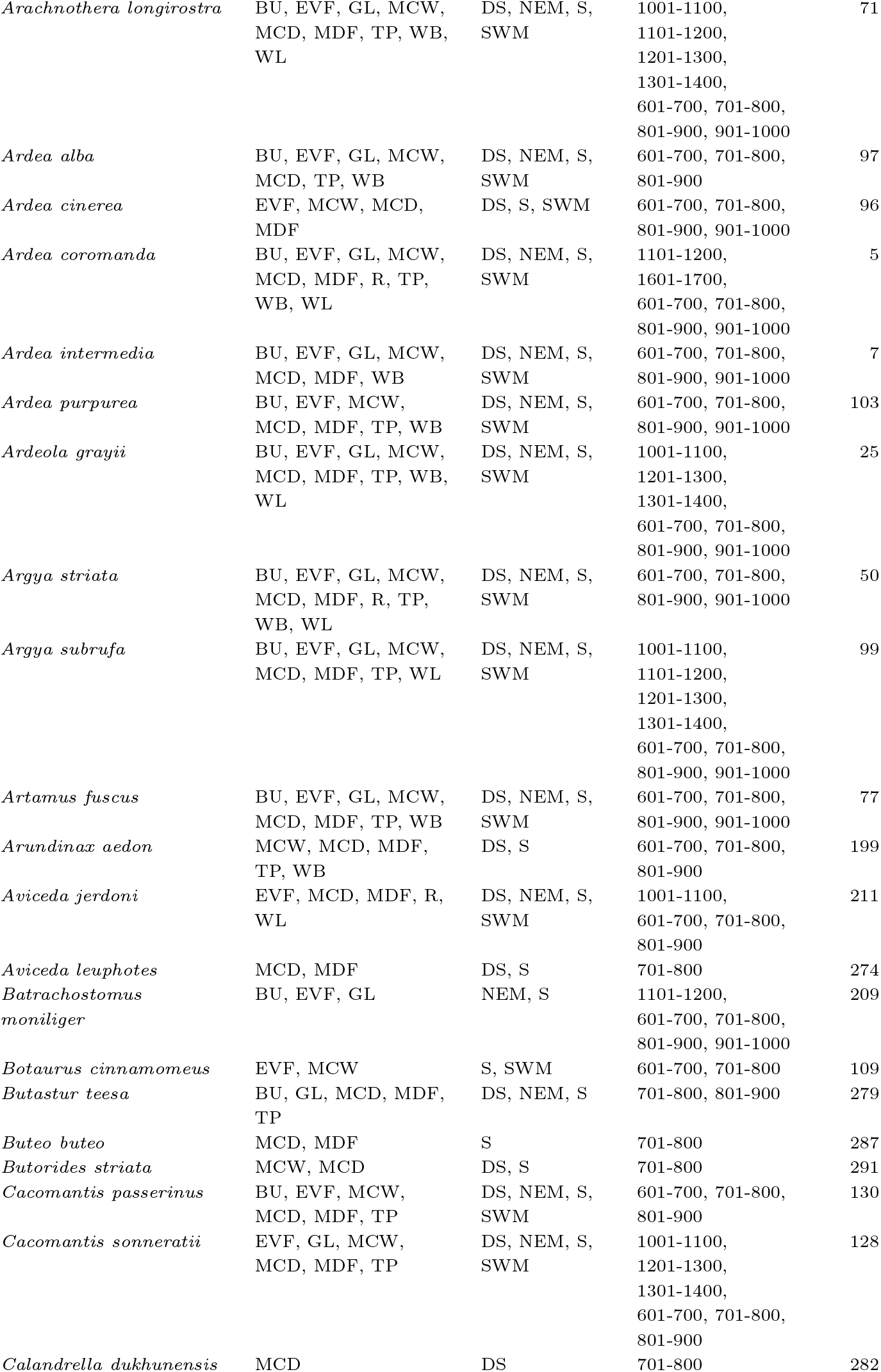
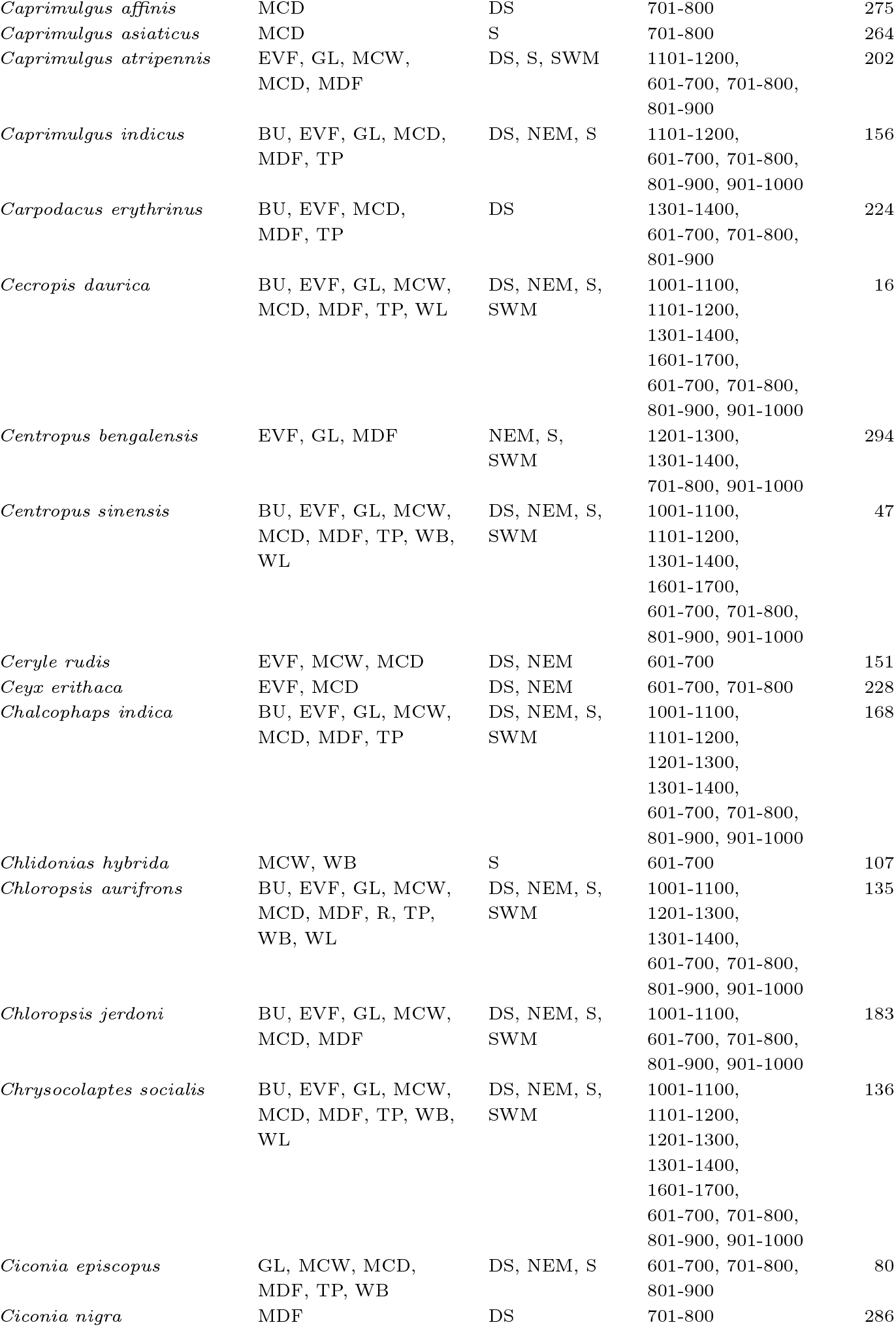
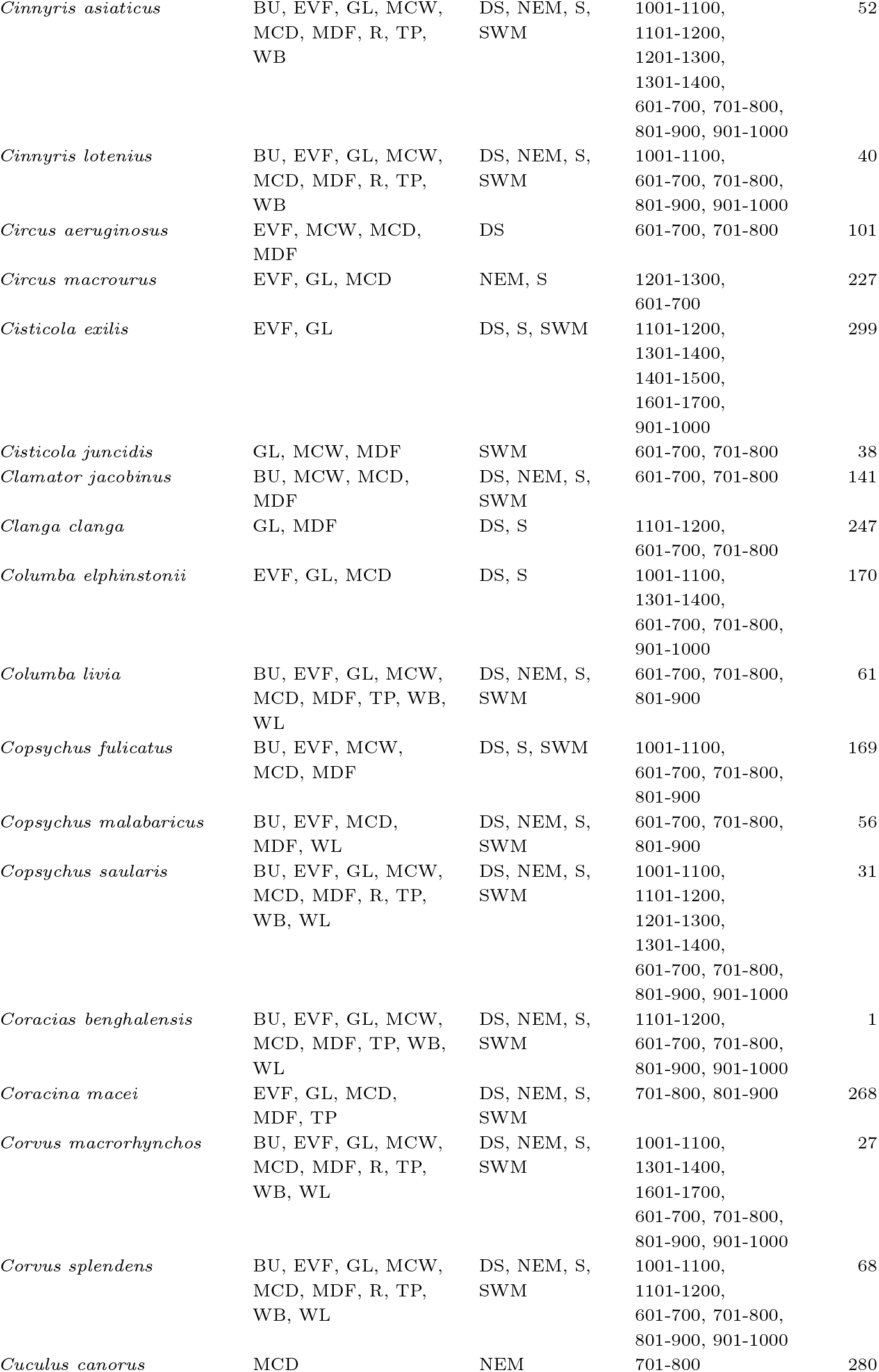
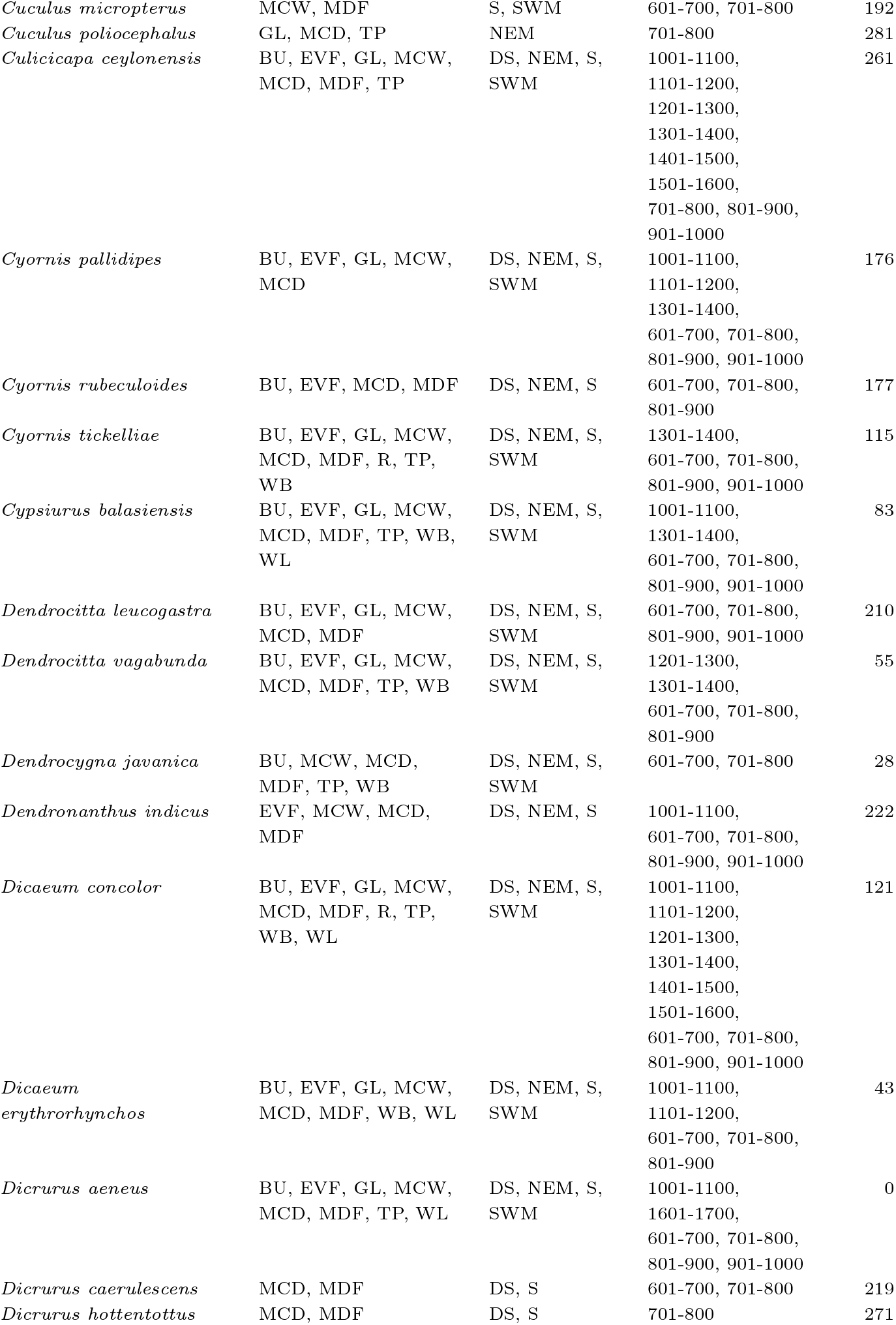
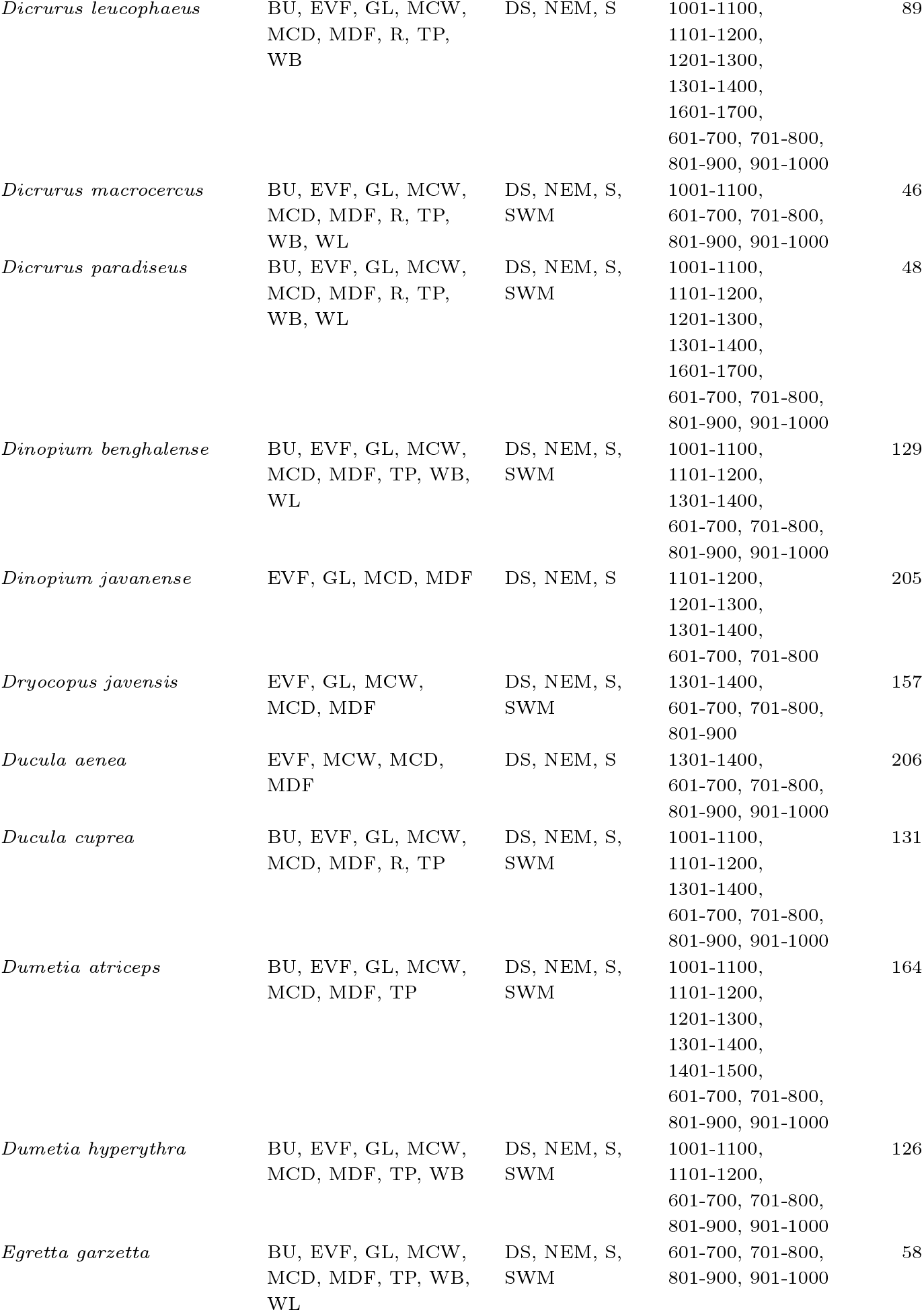
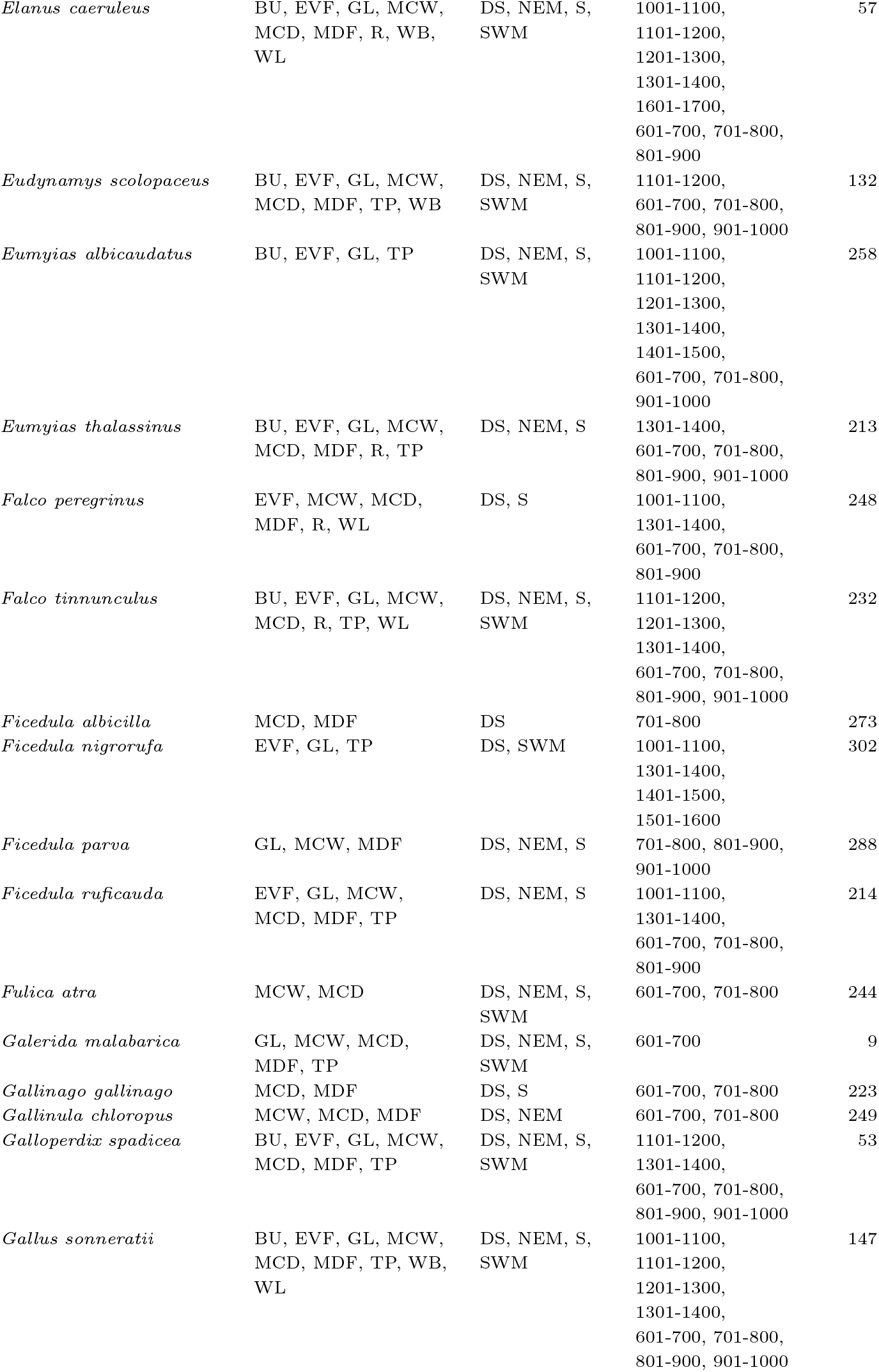
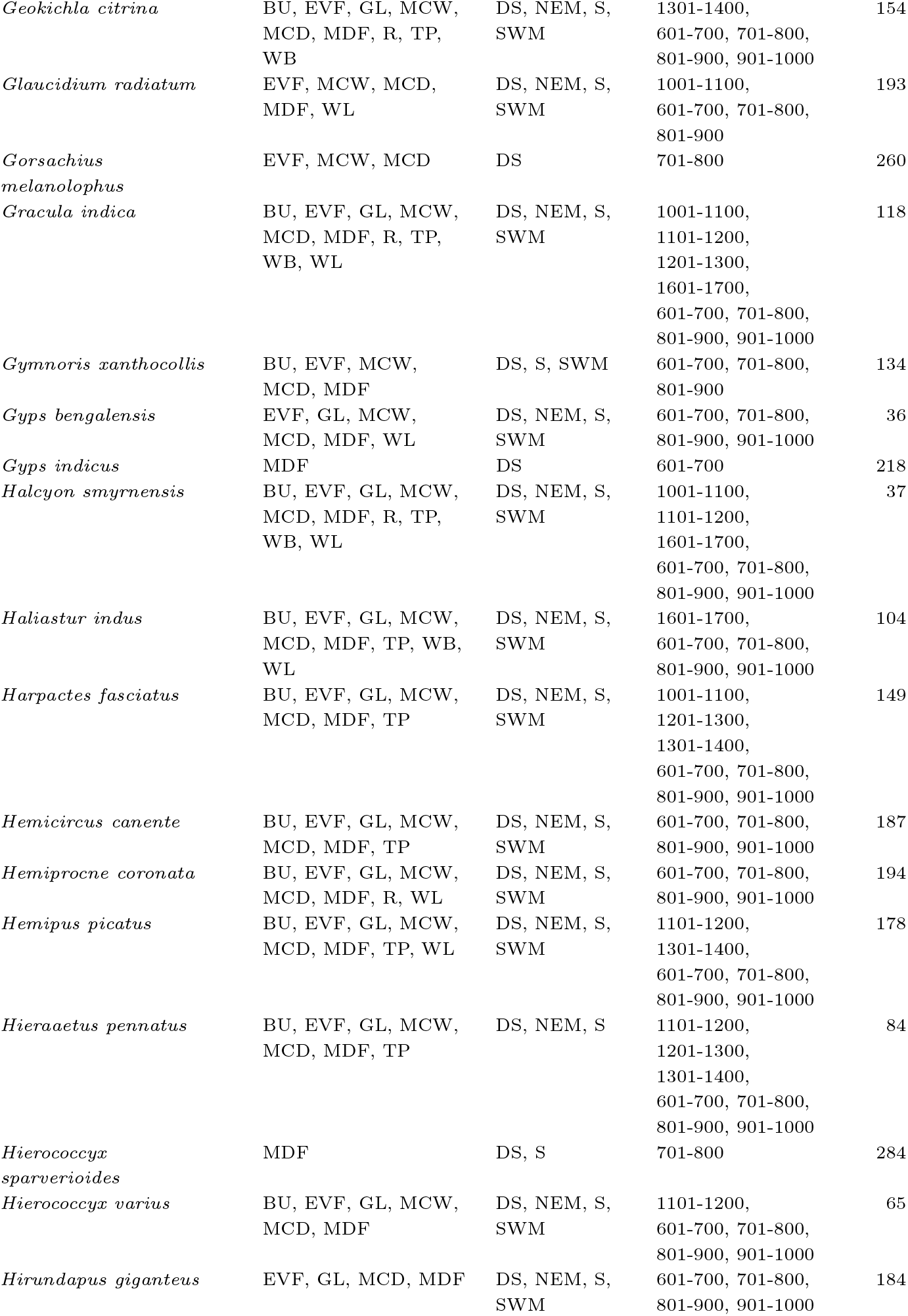
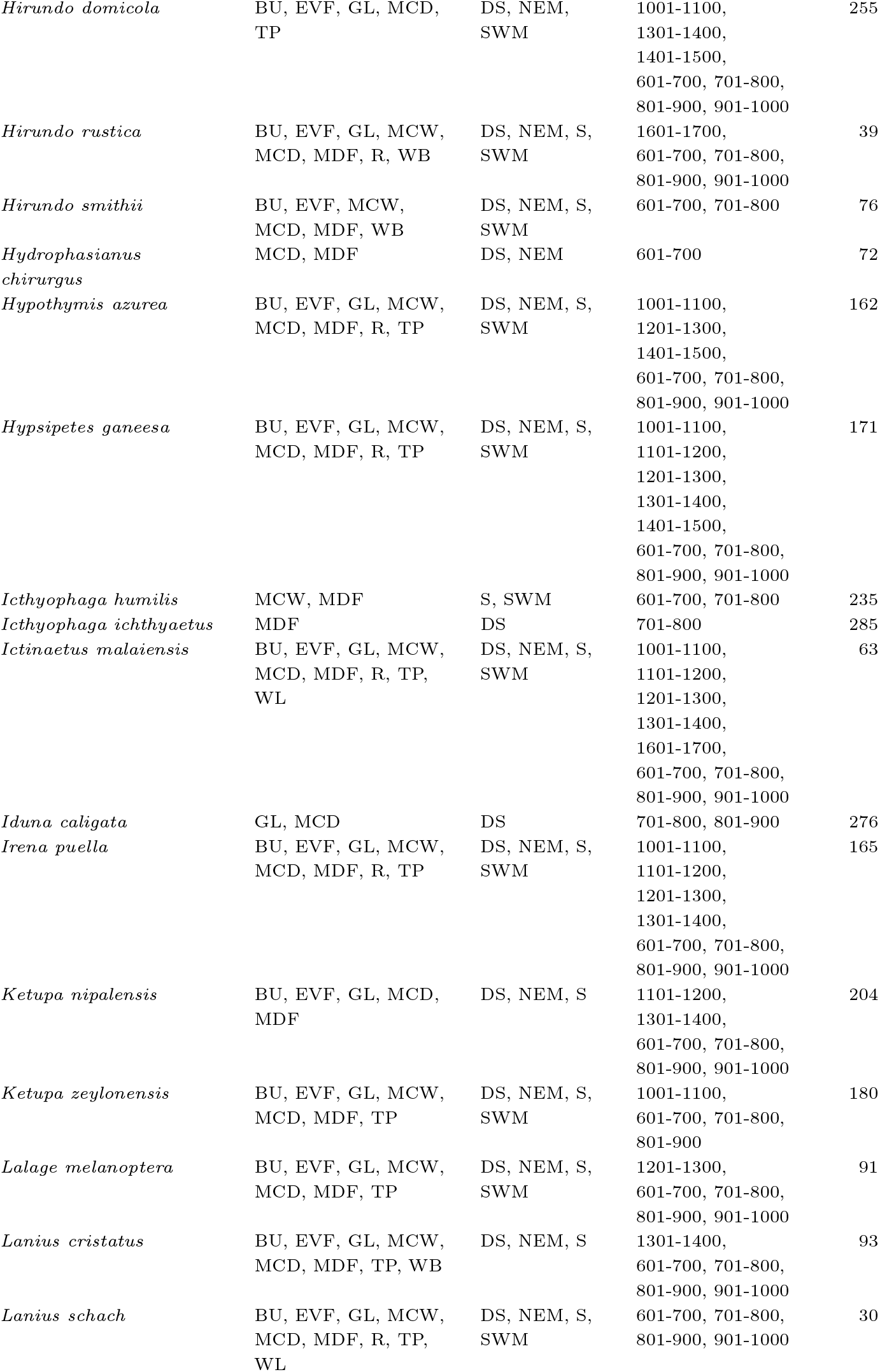
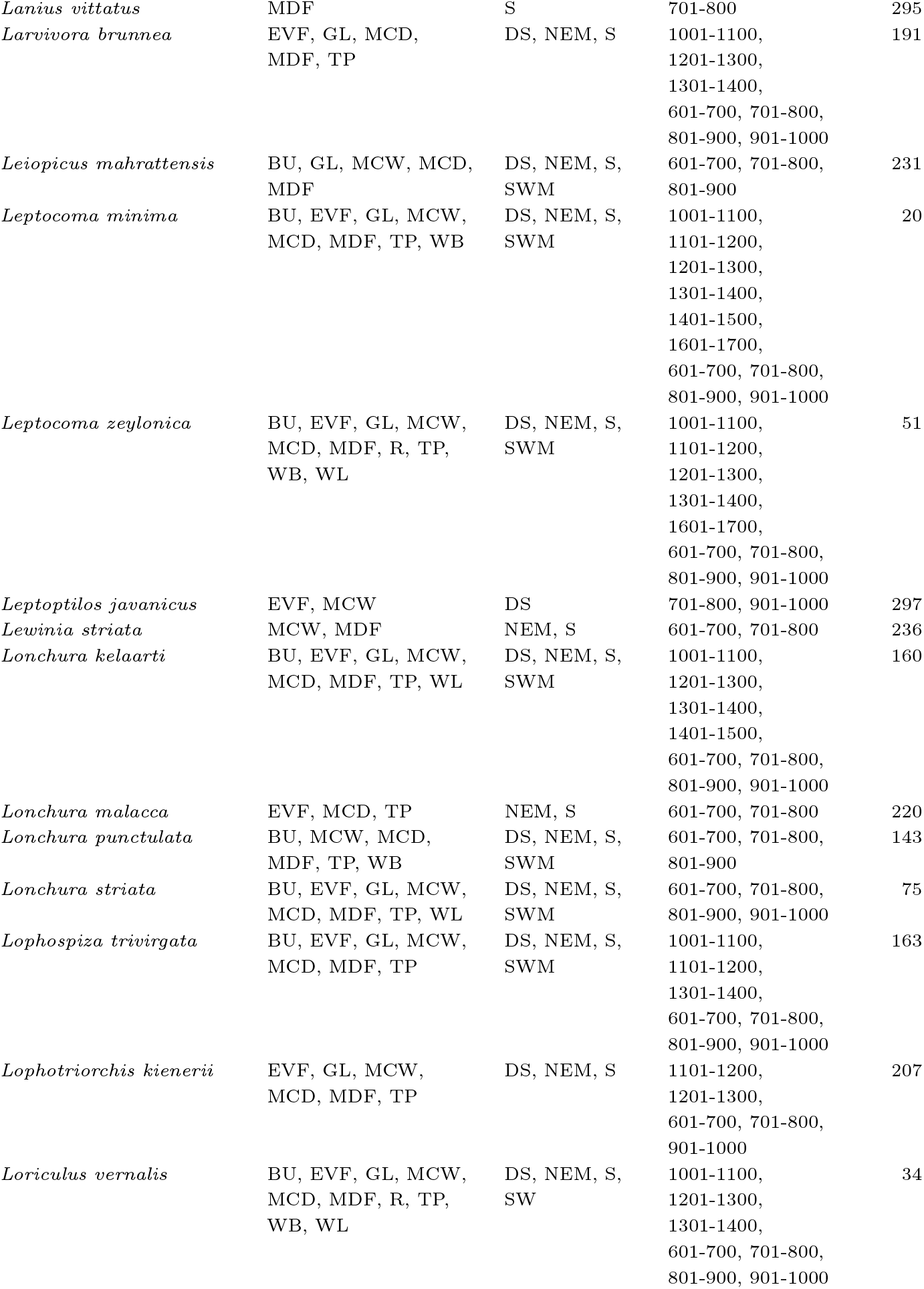
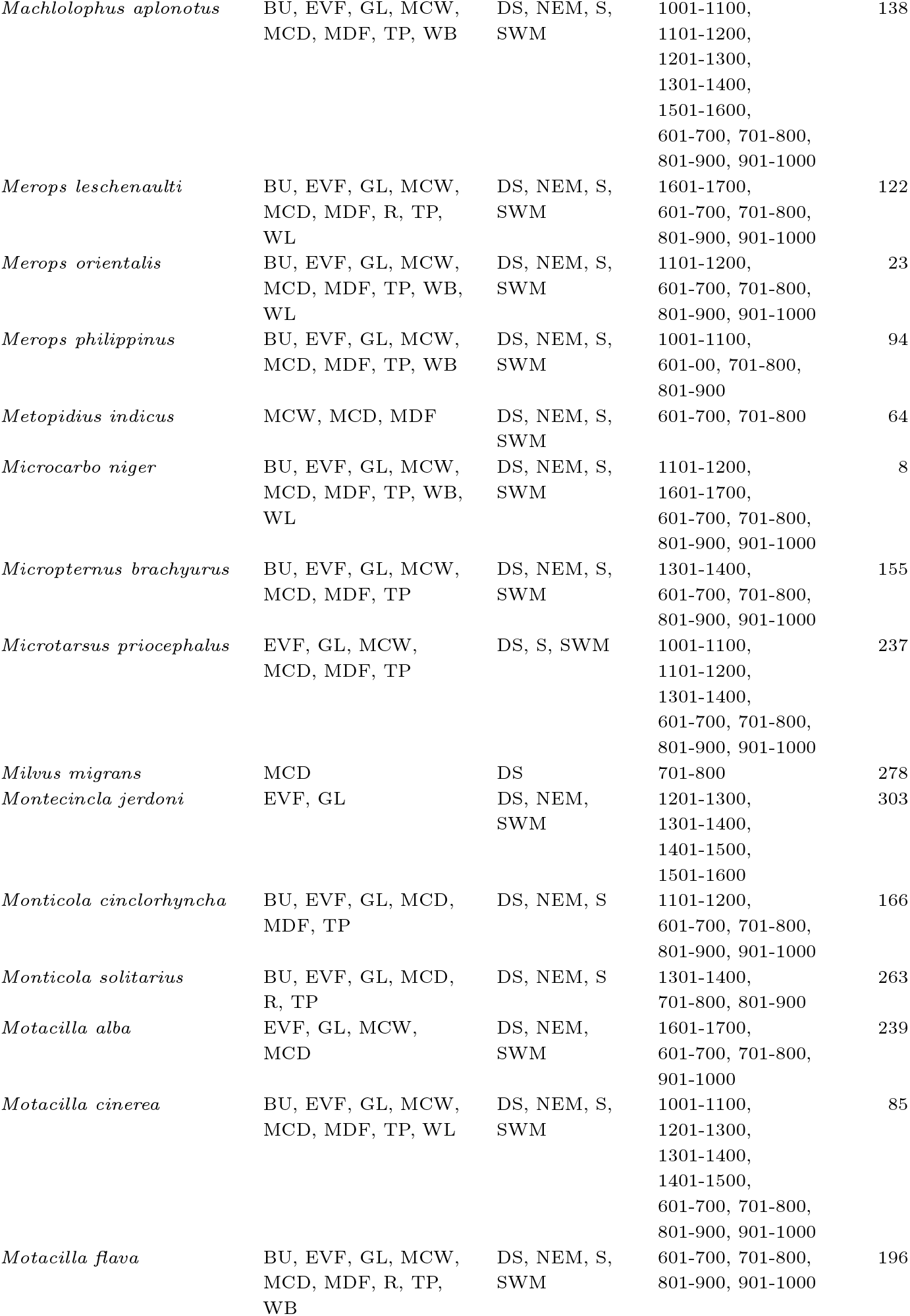
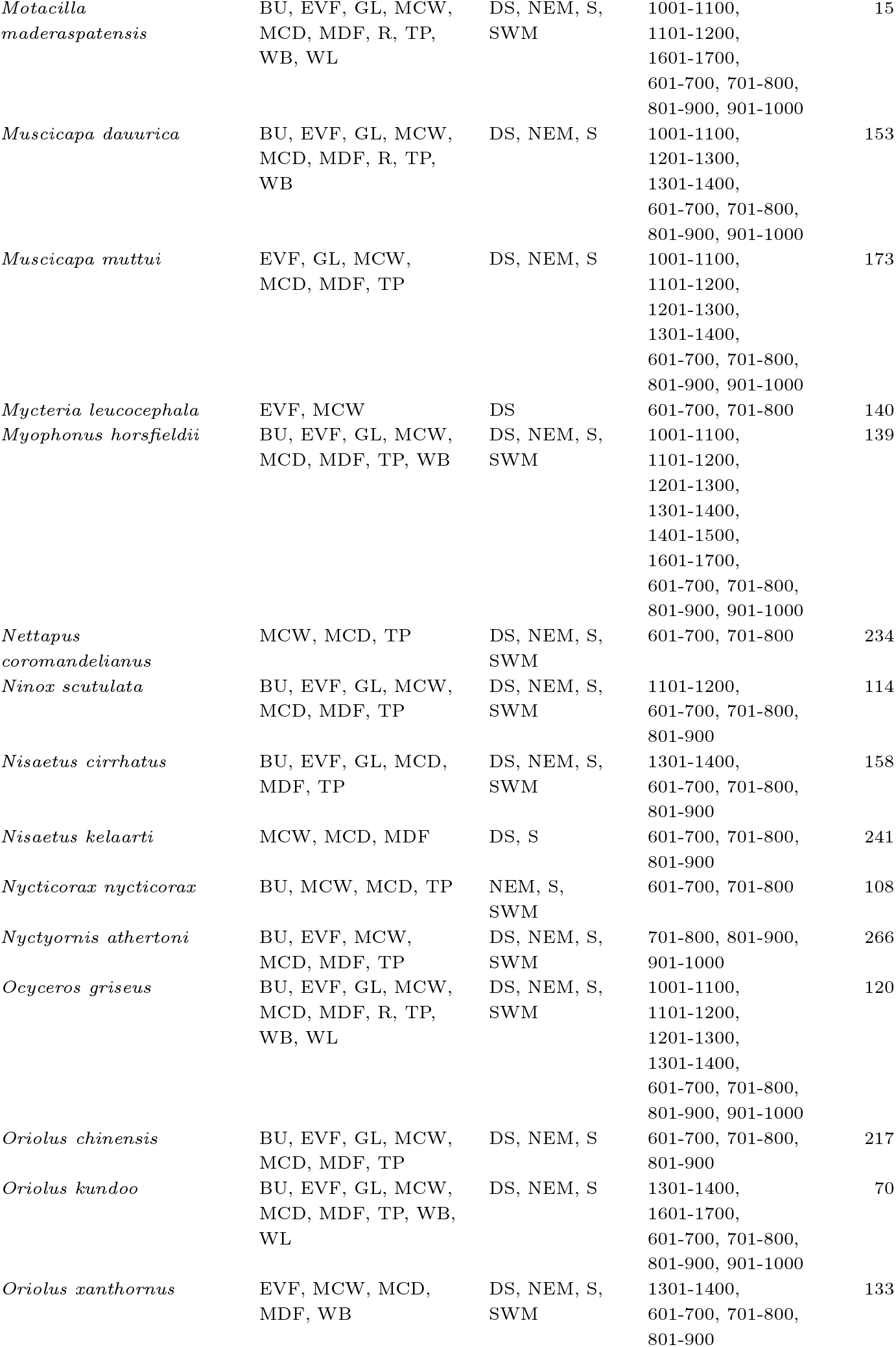
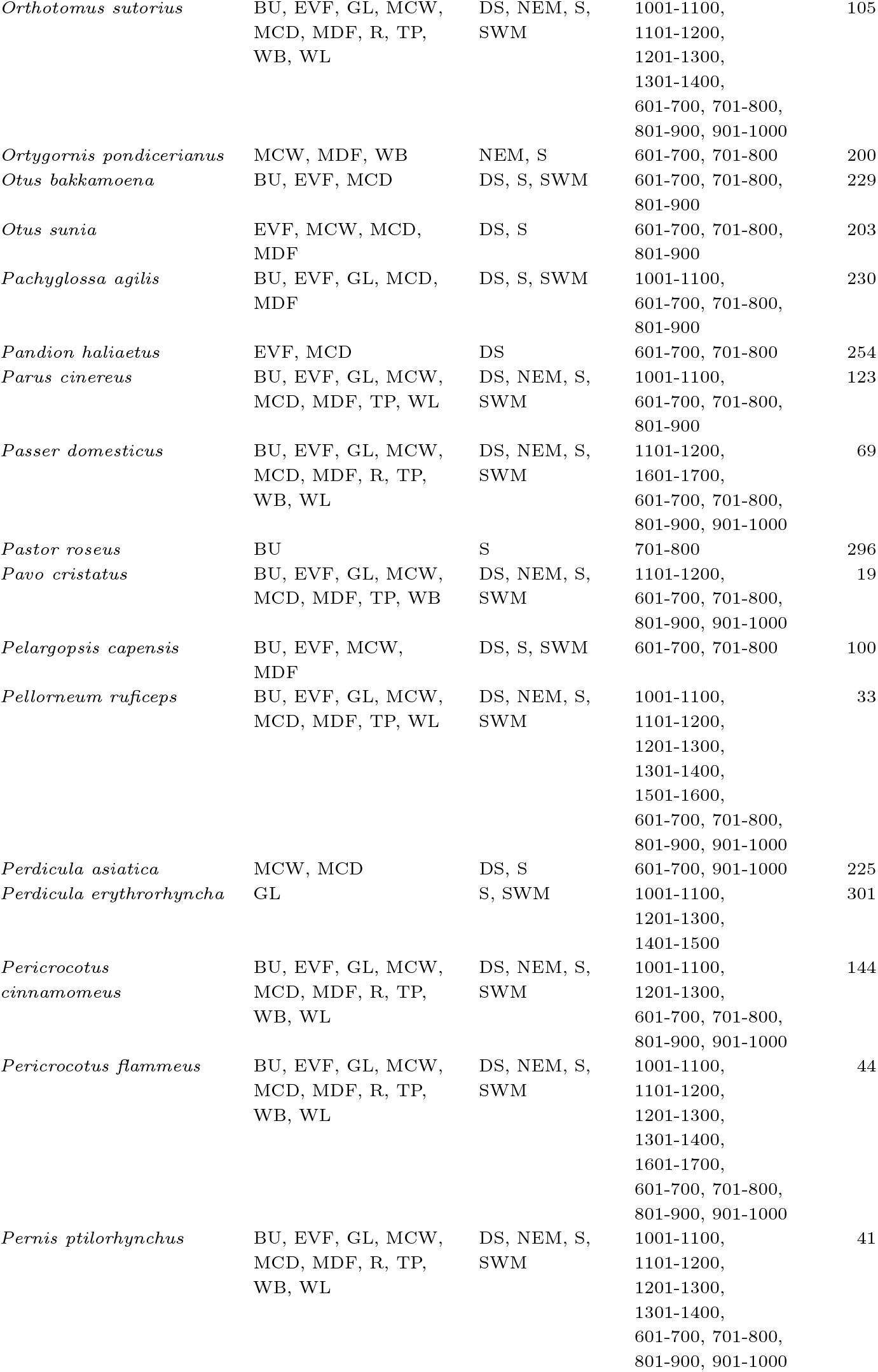
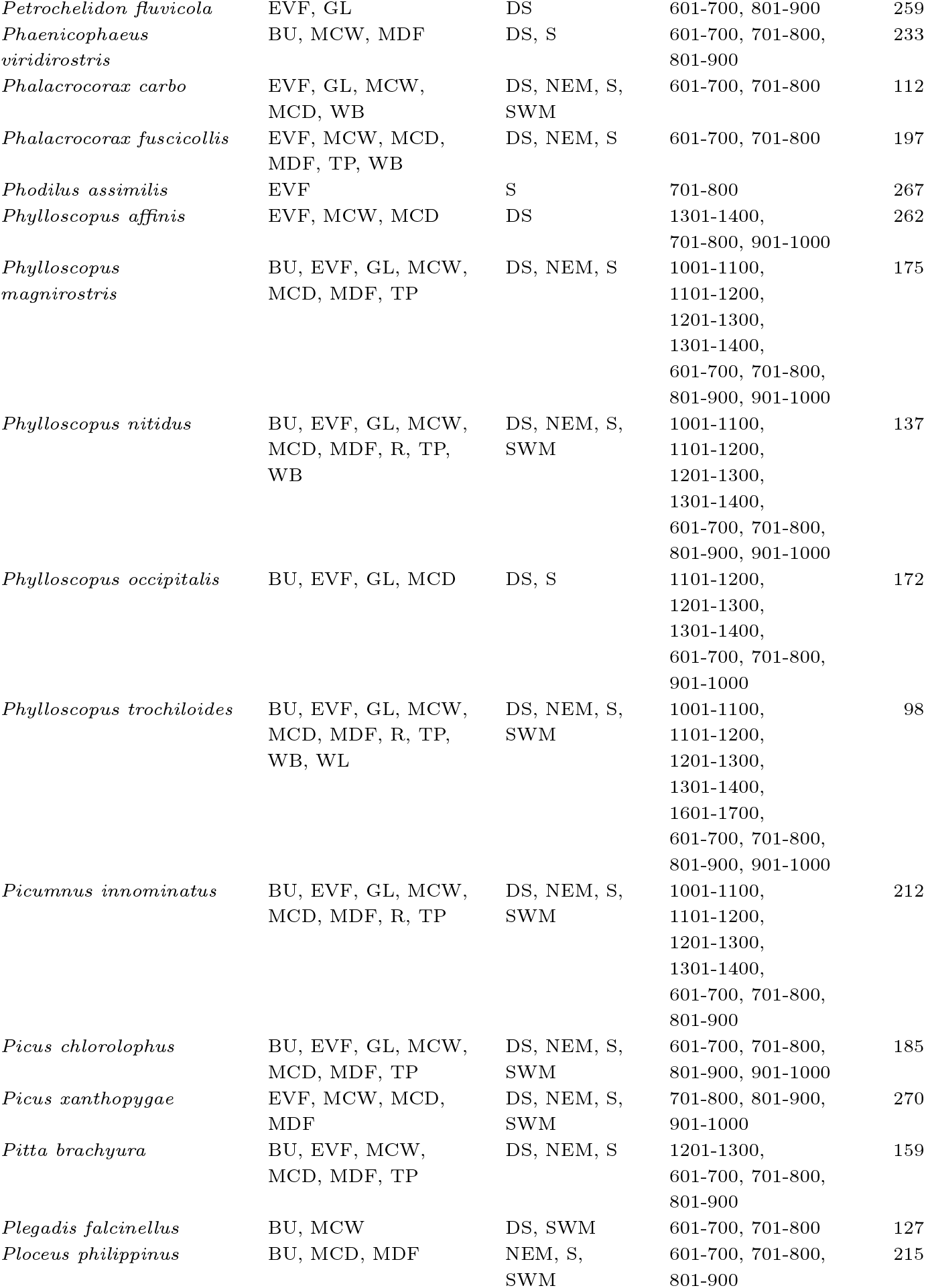
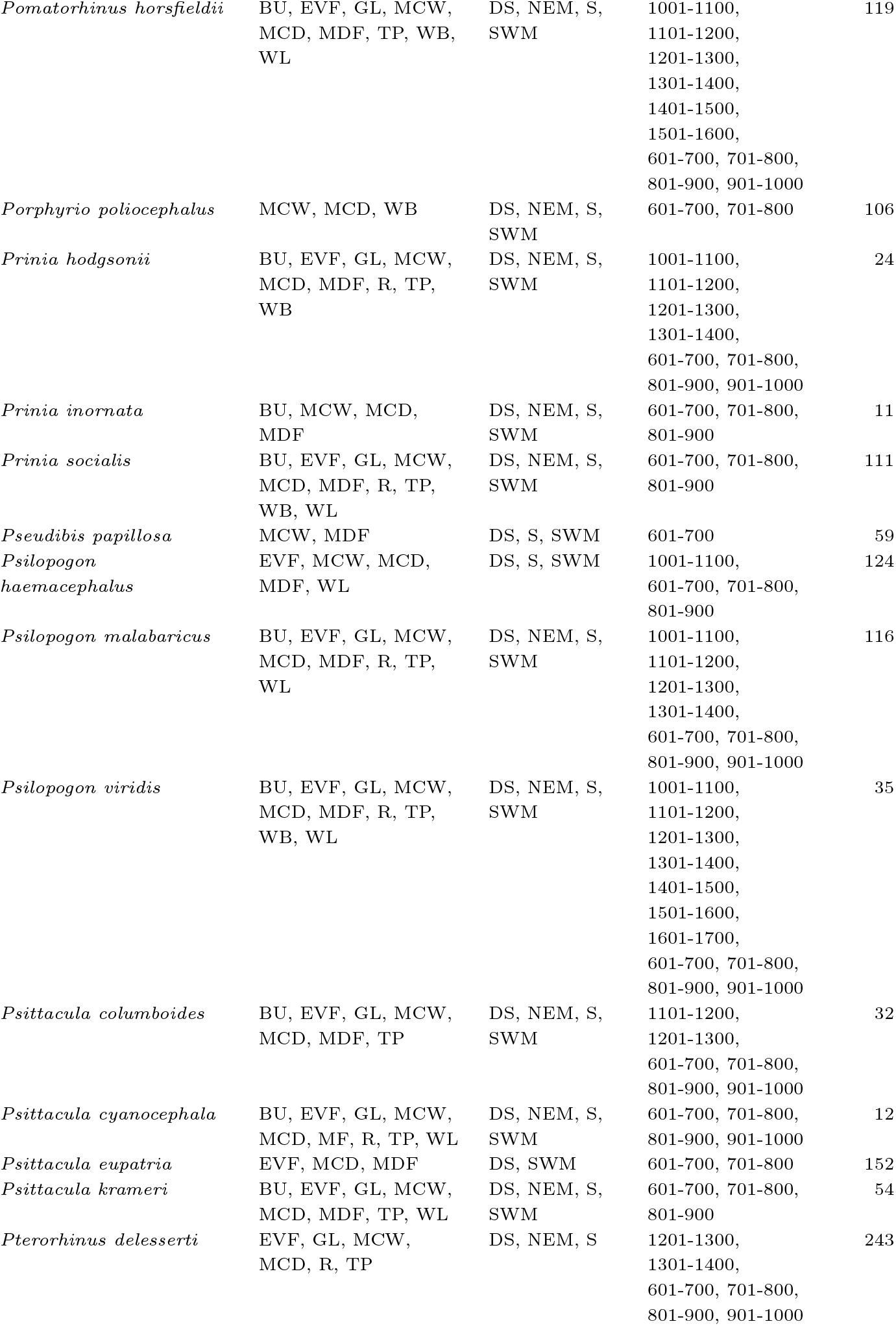
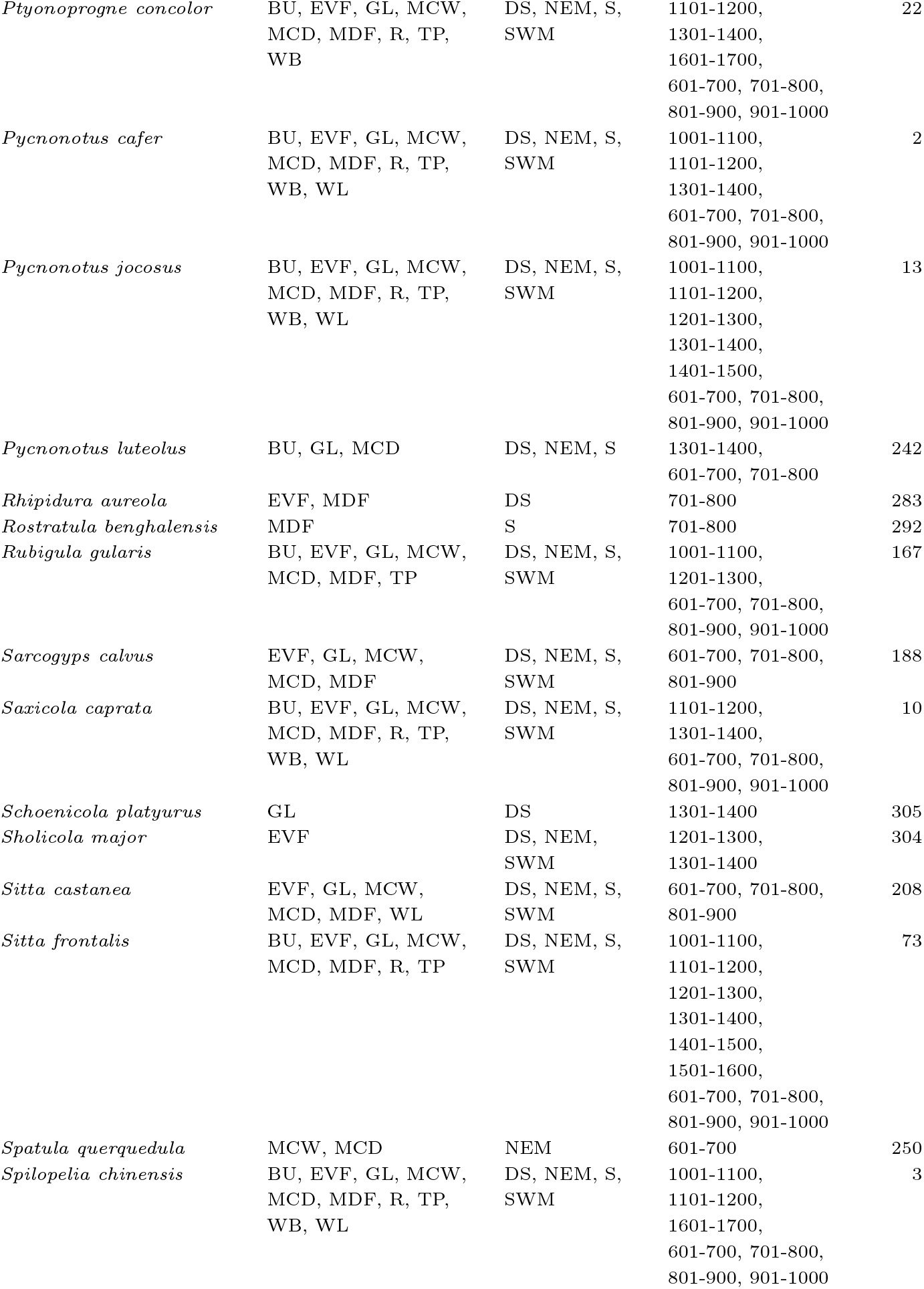
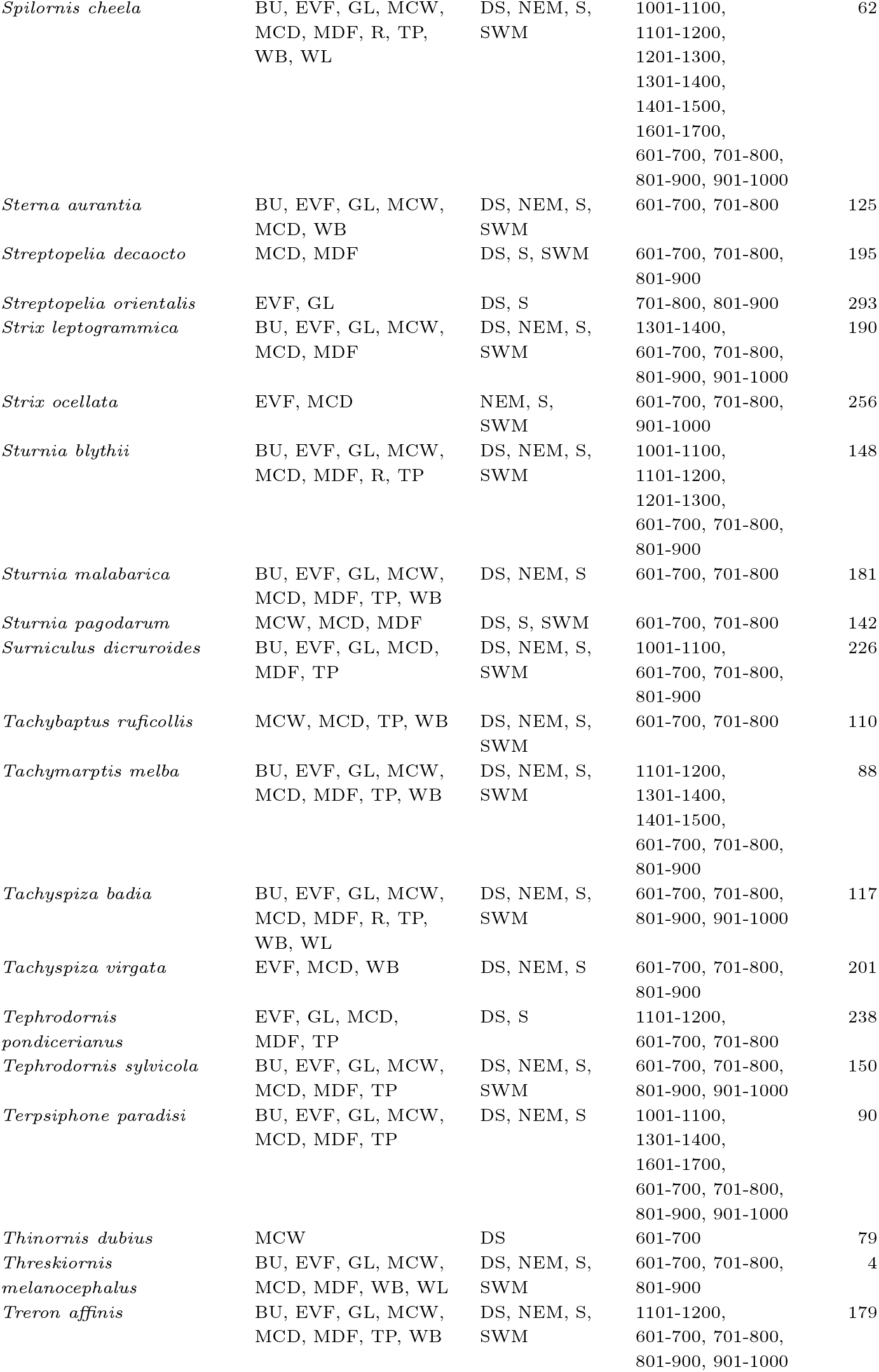
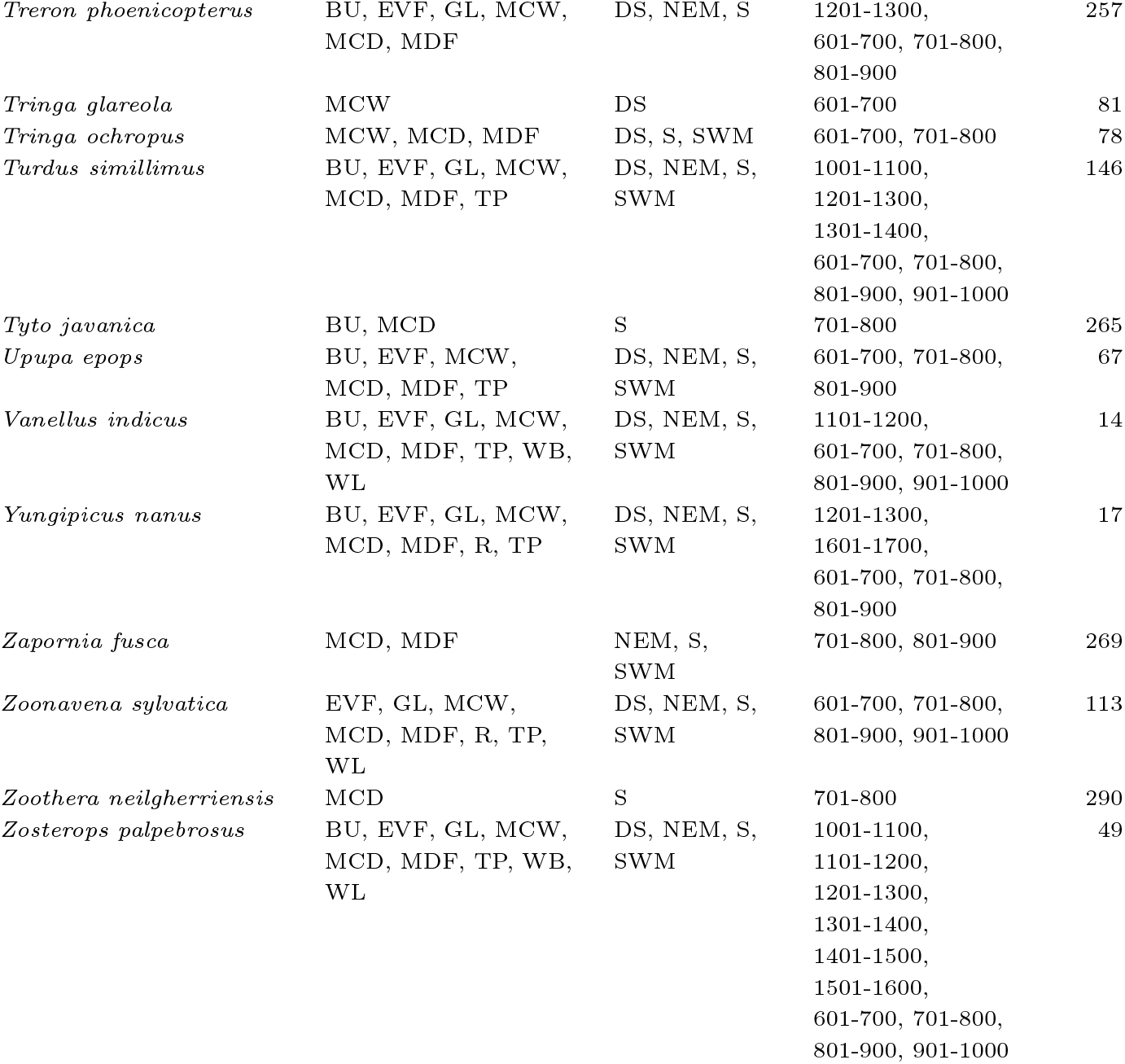
Appendix.

