## Supplementary figures and images for "Functional and Taxonomic Diversity of Avian Communities Across Land-Use Gradients in Wayanad Using eBird Data"

### Lulc Chart.jpg

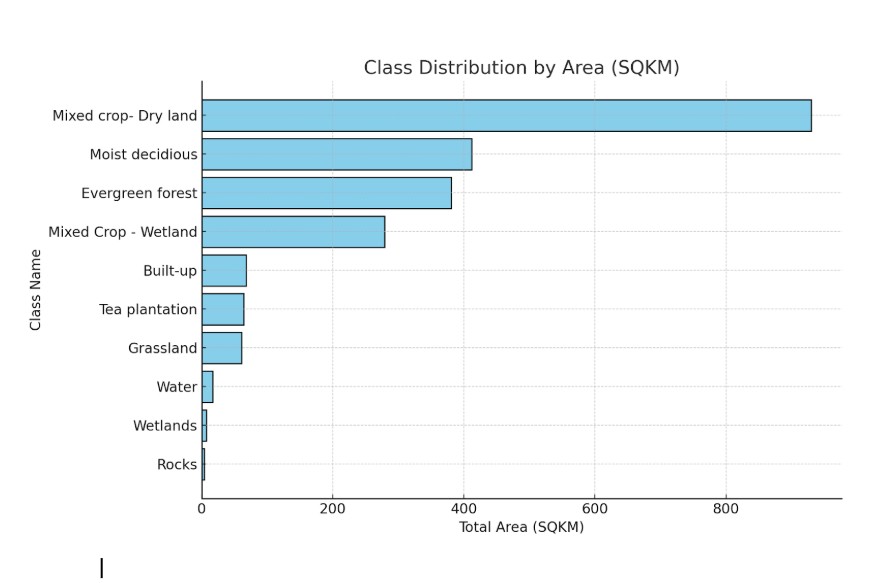
